# Tracking Without Linking or Localization: Single-Photon Tracking

**DOI:** 10.1101/2025.01.10.632389

**Authors:** Lance W.Q. Xu, Nathan Ronceray, Marianna Fanouria Mitsioni, Titouan Brossy, David Šťastný, Jay C. Spendlove, Radek Šachl, Martin Hof, Aleksandra Radenovic, Steve Pressé

**Affiliations:** Center for Biological Physics, Arizona State University, Tempe, AZ, USA; Department of Physics, Arizona State University, Tempe, AZ, USA; Center for Single Molecule Biophysics at the Biodesign Institute, Arizona State University, Tempe, AZ, USA; Max Perutz Labs, University of Vienna, Campus Vienna Biocenter 5, Vienna, Austria; Institute of Bioengineering, School of Engineering, EPFL, Lausanne, Switzerland; NCCR Bio-Inspired Materials, EPFL, Lausanne, Switzerland; J. Heyrovský Institute of Physical Chemistry, Dolejškova 2155/3, CZ-18223 Prague 8, Czech Republic; Faculty of Science, Charles University, Prague, Czech Republic; School of Molecular Sciences, Arizona State University, Tempe, AZ, USA

## Abstract

Mobile biological particles, ranging from biomolecules to viral capsids, often diffuse faster than 1 *µ*m^2^/s, resulting in severe motion blur in conventional millisecond-scale imaging. Shorter exposures capture faster dynamics, but quantizing the per-pixel signal at such exposures eventually degrades our ability to localize particles at all. In the extreme case of binary (1-bit-per-pixel) output—where moving from 8-bit grayscale to 1-bit corresponds to a 255-fold faster acquisition rate—no existing tracking method applies, since all of them rely on intensity-based localization that binary data cannot supply. We therefore introduce single-photon single-particle tracking (SP^2^T), a framework that bypasses localization and linking by estimating particle numbers and trajectories directly from the full 1-bit image stack. We demonstrate SP^2^T on single-photon avalanche diode (SPAD) arrays, which deliver microsecond exposures over a large field of view (512×512 pixels) and explicitly account for detector-specific artifacts such as hot and cold pixels. We validate the method against programmed fluorescent bead and organic dye trajectories, and apply it to two biological systems, aerolysin and ganglioside GM1. Together with simulation, these experiments reveal that longer exposures bias diffusion coefficient estimates (by up to 70% for particles diffusing at 5 *µ*m^2^/s) and distort jump-distance distributions, underscoring the need for photon-by-photon tracking in fast-diffusion regimes. Evaluating the likelihood on a GPU further delivers more than a 50-fold speedup over a CPU implementation when compared on datasets of the same frame size and number of frames, enabling multiplexed tracking for crowded particles. Together, these advances establish SP^2^T as a robust, data-efficient route to unbiased particle tracking at microsecond temporal resolution.

## Introduction

Advances in fluorophore photophysics have allowed us to overcome the diffraction limit of light, enabling fluorescent localization of molecular positions with remarkable precision, down to approximately 70 nm(*1–14*). More recent techniques have pushed this precision even further, with localization down to 6 nm achieved through excitation beam modulation(*15*), and subsequent *in vitro* techniques reaching sub-nanometer precision(*16*). With these advancements, we can now directly observe several biological structures(*17*), nanoparticle endocytosis, and targeted drug delivery to cancer sites(*18, 19*), as well as viral particle activity within living systems(*20, 21*).

Despite remarkable advances in spatial resolution, achieving high temporal resolution remains challenging. Motion blur arises readily at typical exposure times (≈10 ms), with visible artifacts even at diffusion coefficients as low as 1 *µ*m^2^/s (Fig. 1b). This limitation becomes critical when imaging faster diffusing molecules: SARS-CoV-2 virions diffuse at rates exceeding 5 *µ*m^2^/s(*22*), and cytoplasmic proteins can diffuse faster than 10 *µ*m^2^/s(*23*). Moreover, motion during exposure not only degrades localization precision but also introduces ambiguity in identifying individual molecules, especially when blurred signals overlap with nearby emitters. Recent work with MINFLUX(*15*), a real-time tracking method, shows that slow acquisition rates may even systematically underestimate diffusion coefficients(*24*), underscoring that acquisition speed remains a fundamental constraint across modalities. These challenges are further exacerbated by the diffraction-limited point spread function (PSF), which causes even small biomolecules to appear as extended features (about 250 nm)(*25*).

**Figure 1:**
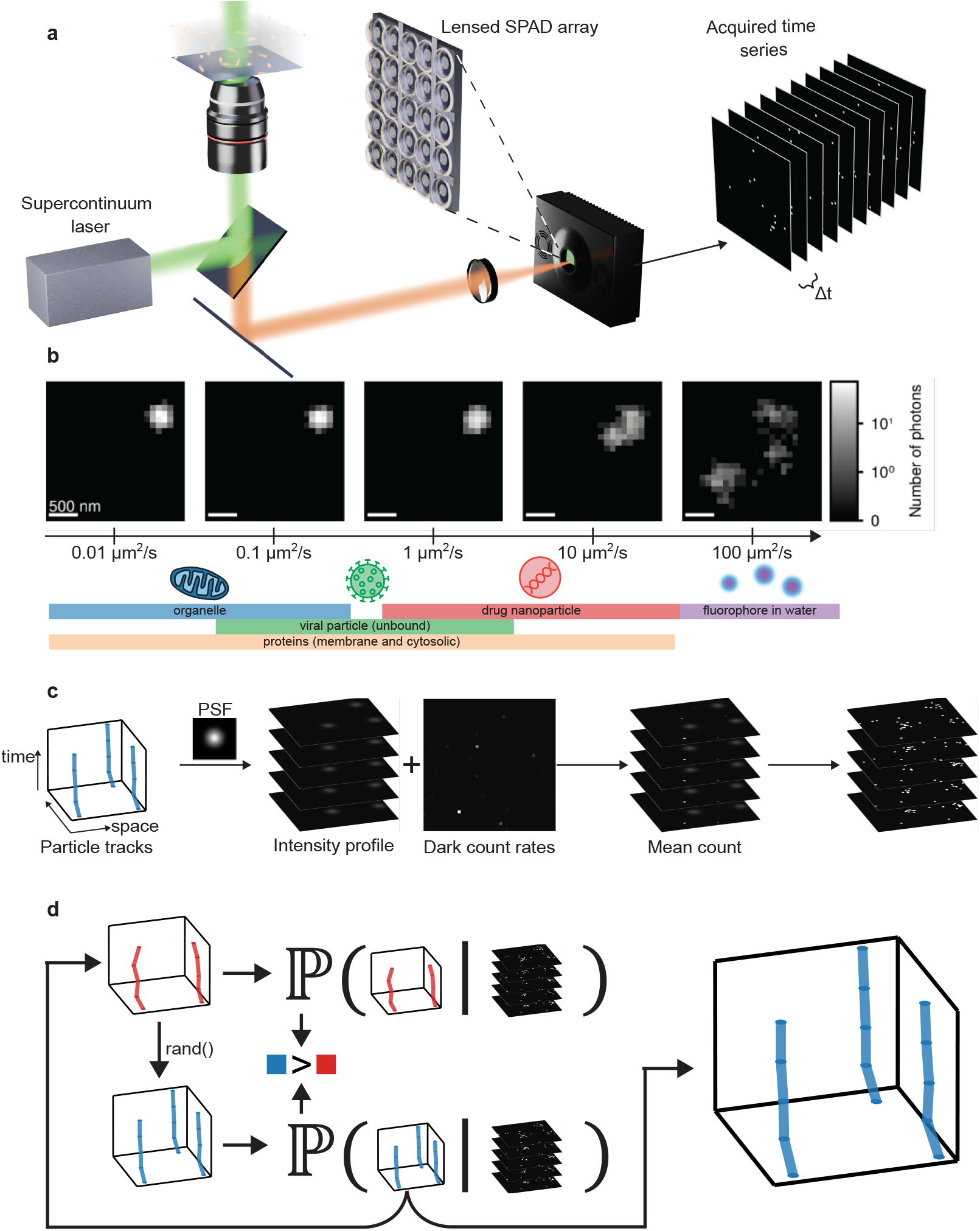
**a**, Schematic of the experimental setup. A supercontinuum pulsed laser source provides illumination, with the option to switch between epi-illumination and total internal reflection fluorescence modes. Image acquisition is performed using the SPAD512^2^ camera at frame rates up to 97.7 kHz. Only a subset of the full 512*×*512 pixel field of view (FOV) is shown for illustrative purposes. **b**, Typical diffusion coefficient ranges for different biological systems(*49*). Motion blur begins to appear at diffusion coefficients as low as 1 *µ*m^2^/s with an exposure time of 10 ms, potentially causing a single particle to be misinterpreted as multiple distinct objects, as shown in Fig. S.2 and as becomes apparent as we move beyond 1 *µ*m^2^/s. **c**, As a physics-inspired model, SP^2^T considers the process by which data is generated from the fluorescently labeled particles, including the system’s point spread function (PSF) and the dark count rate of each pixel—spurious detections registered in the absence of incident light, which vary from pixel to pixel and are calibrated per detector. **d**, A Monte Carlo algorithm to sample candidate particle numbers and tracks from the posterior probability distribution. Based on the current track sample (red), we propose a new particle with its associated track (blue) and compute corresponding posterior probabilities. Favoring higher probability, we stochastically accept or reject this new track and update the current sample. In this way, we propose particle numbers and their associated tracks consistent with the entirety of the FOV over all frames and all of the optics systematically integrated into the probability.

Stroboscopic illumination partially mitigates motion blur by narrowing the effective exposure window(*26–28*). However, by sampling intermittently, it risks missing transient events between flashes, introducing tracking errors such as trajectory mislinking (Fig. S.1). As a result, while effective for reducing blur, stroboscopic imaging provides only a partial solution to the problem of temporal resolution. Recently, a single-objective light-sheet variant (oblique line-scan illumination) demonstrated the tracking of very fast-diffusing proteins (up to about 14 *µ*m^2^/s in living cells)(*29*), yet it still inherits the intermittent-sampling issue characteristic of stroboscopic schemes.

To truly overcome these limitations, continuous high-speed acquisition is required. Cameras capable of microsecond-scale exposures, typically based on scientific CMOS sensors or single-photon detectors (SPDs), are increasingly central to this effort. SPDs encompass several device classes, including photomultiplier tubes, single-photon avalanche diodes (SPADs), superconducting nanowire single-photon detectors, and transition-edge sensors(*30*). These detectors are increasingly adopted in biological imaging because they have no readout noise and can perform parallel single-photon counting and time-resolved measurements(*31–38*).

In this work, we focus on SPAD arrays, one detector architecture that produces rapid single-photon-per-pixel output, and specifically on the SPAD512^2^ from Pi Imaging. This sensor combines a large field of view (512*×*512 pixels) with a high data acquisition rate of 97.7 kHz, making it well-suited for high-speed imaging applications. Designed to operate reliably at or slightly below room temperature, the SPAD512^2^ features a compact form factor, low operating voltage, and robust performance across a range of conditions(*39*). SPAD arrays have already demonstrated utility in diverse domains, including fluorescence microscopy(*31–33, 35, 36*), time-resolved spectroscopy(*40–42*), and nuclear medicine(*42*). Beyond their exceptional temporal resolution, SPAD technology also enables fluorescence lifetime imaging, providing detailed insights into molecular species and their local environments.

Despite the rapid data acquisition rates of SPAD arrays and other devices capable of raw 1-bit (single-photon) output, no existing method can directly extract dynamical information from such data. The central challenge is that tracking requires localizing light-emitting particles in each frame, yet most localization algorithms depend on sufficient detected photons (*i*.*e*., pixel intensities) to achieve high accuracy, as quantified by the Cramér-Rao lower bound(*43, 44*). This requirement fundamentally conflicts with the discrete, single-photon nature of SPD data. Consequently, recent studies with SPDs, including SPAD arrays, still rely on binning consecutive 1-bit frames for analysis(*38, 45*), a process that sacrifices the exceptional temporal resolution these detectors can provide (Fig. 1). While binning enables particle localization with conventional tracking tools, it does not add information; rather, it averages away temporal information contained within the dataset(*46–48*). This highlights the need for methods that bypass binning and instead leverage the full information content of raw 1-bit frames.

Here, we introduce single-photon single-particle tracking (SP^2^T), a framework designed to directly track particles from the fastest acquirable 1-bit frames. Rather than localizing particles independently in each frame and subsequently linking those localizations into trajectories, SP^2^T jointly infers particle numbers and trajectories in a manner consistent with the entire 1-bit-per-pixel image stack. Extracting dynamics at this temporal resolution, however, requires careful calibration of pixel-specific detector properties. This is particularly important for SPAD arrays, where pixel-to-pixel variations are more consequential than in integrative detectors, whose analyses are generally more robust to such variations in noise properties(*25*). By explicitly accounting for these detector-specific effects, SP^2^T avoids mistaking hot pixels (which fire far more often than the incident light warrants) for particles or interpreting cold pixels (which rarely fire at all) as regions systematically avoided by particles. Together, these features allow SP^2^T to bypass the conventional frame-by-frame localization and linking paradigm and push particle tracking toward the fastest timescales accessible to single-photon detectors. In doing so, we highlight the consequences of longer exposures for diffusion coefficient estimation and diffusive-model identification, in diffusing ganglioside GM1 molecules and in simulated fast-diffusing particles.

## Results

To validate that SP^2^T can reliably track particles from 1-bit frames without either localizing in each frame or linking localizations frame-to-frame, we begin by fixing fluorescent beads to a stage programmed to follow a known sequence of steps, which lets us compare inferred tracks both against the known stage motion and against one another. We then use surface-immobilized Atto643 organic dyes also mounted on a moving stage to test the robustness of tracking under dimmer conditions.

Next, using a dataset of diffusing ganglioside GM1 molecules, we show how analyzing at higher temporal resolution changes diffusion coefficient estimates. We then highlight the multiplexing capability of our tool by analyzing an aerolysin dataset, tracking monomers in bilayers over a large imaging region. To further demonstrate the multiplexing ability of SP^2^T to simultaneously track many emitters across a large FOV at single-photon frame time resolution, we track 100 particles at 100 *µ*s exposures using synthetic data where ground truth is known, mimicking the aerolysin data. Finally, we assess the robustness of SP^2^T in reliably tracking closely spaced emitters using pseudo-synthetic data. Additional synthetic data robustness tests are included in the Supplementary Information, including robustness analyses on diffusion coefficient (Fig. S.3), emission brightness (Fig. S.4), and dark count rates (Fig. S.5).

As SP^2^T is based on a probabilistic framework within a nonparametric Bayesian paradigm(*50*), it returns full posterior distributions over particle numbers and associated tracks rather than point estimates. For legibility, we plot the *maximum a posteriori* (MAP) sample, and quantify uncertainty from ∼500 posterior samples per track.

### Tracking emitters on a moving stage

The first control experiments we present involve fluorescent beads fixed to a stage following a pre-programmed trajectory. The trajectory alternates between holding the stage stationary for a fixed dwell time (40 ms) and moving it by a predetermined step size (∼170 nm) in a randomly chosen direction: left, right, up, or down.

This setup allows us to: 1) qualitatively compare our tracking results against the programmed sequence of steps; 2) assess the consistency of the inferred tracks across multiple beads, thereby confirming the robustness of our framework; and 3) benchmark our results against those obtained using TrackMate(*51*) on binned frames. These comparisons demonstrate that SP^2^T resolves particle tracks with sub-pixel accuracy while achieving at least an order of magnitude finer temporal resolution.

In Fig. 2a, we first plot the inferred particle tracks on top of the sum of all binary frames. Over the approximately 115 ms dataset, the tracks clearly reproduce the three expected programmed ∼170 nm stage movements consistent with experimental settings— left, up, then down. These shifts occur at approximately 15 ms, 55 ms, and 95 ms, spaced by the expected dwell time of 40 ms. We then align the three tracks by subtracting each track’s center of mass, revealing strong sub-pixel agreement. During the stationary dwell periods at the vertices of the “L”-shaped trajectory, the inferred positions are confined within approximately 50 nm. Across the full trajectories, the average pairwise distance between aligned tracks is approximately 21 nm, while the maximum separation remains below 65 nm, all substantially smaller than the 100 nm pixel width and 230 nm PSF radius (fitted), despite being reconstructed from 5,750 1-bit frames with about five photons per bead per frame. This agreement is shown in greater detail in Figs. 2b and 2c, where the *x*- and *y*-tracks are overlaid on the binned marginal posterior density (confidence) of bead 1. The spread of this posterior provides a measure of localization uncertainty, with an average standard deviation of approximately 16 nm for each bead.

**Figure 2:**
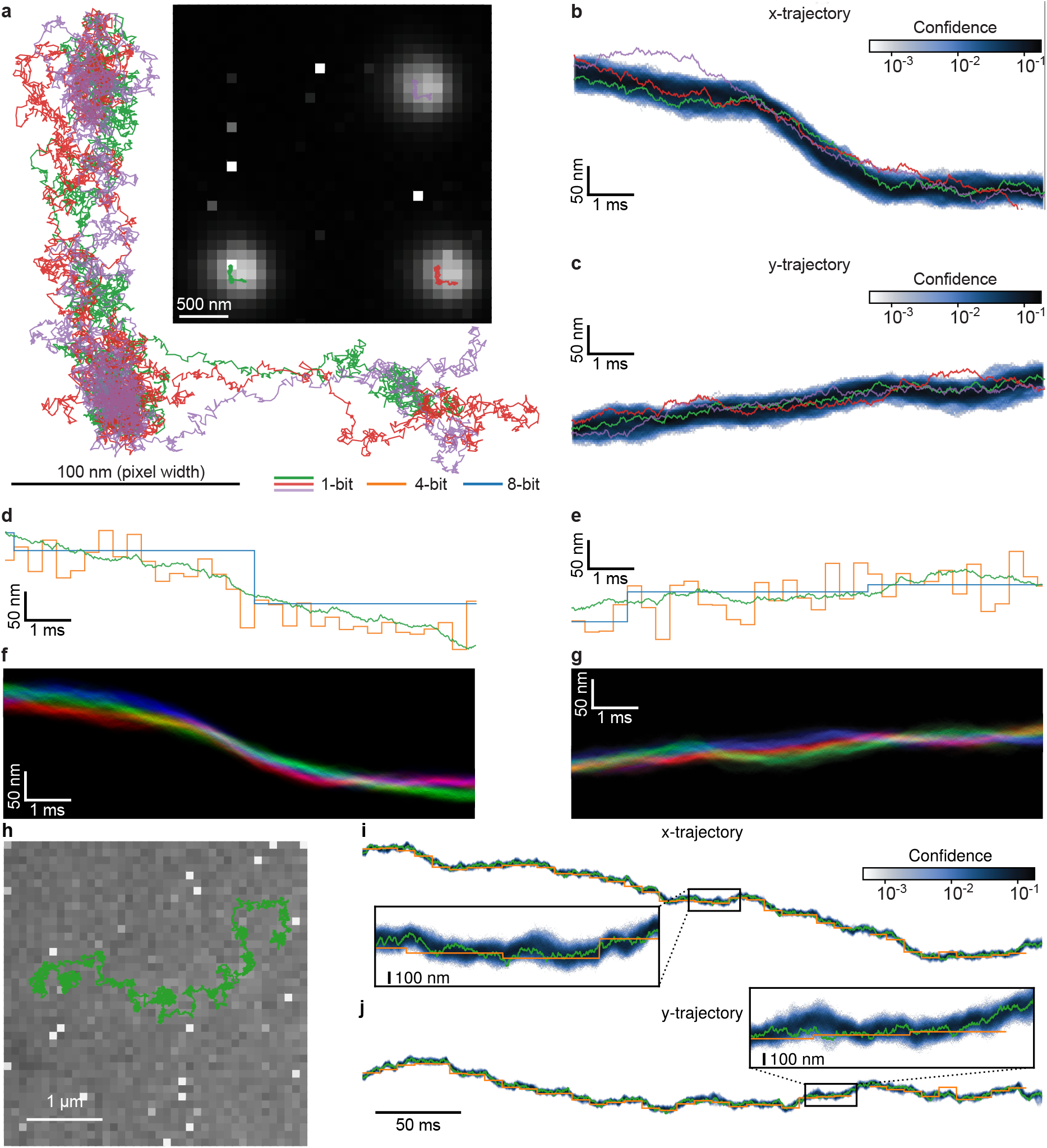
Results of broadband fluorescent beads fixed to a moving stage with 40 ms dwell time and 170 nm step size. Experimental parameters include: numerical aperture of 1.49, refractive index of 1.52, and pixel width of 100 nm, corresponding to a diffraction limit of approximately 230 nm. **a**, Inferred trajectories plotted over the sum of all 1-bit frames with 10 *µ*s exposure, cropped and contrast-adjusted for illustrative purposes. Additionally, the three inferred bead tracks are aligned and overlaid to assess consistency. **b**, Full confidence of bead 1’s *x*-track plotted as a shaded region in a time interval (10 ms to 20 ms); darker colors indicate higher confidence. **c**, The same layout as **b** but for bead 1’s *y*-track. **d**, Direct comparison of bead 1’s 1-bit *x*-track with TrackMate’s *x*-tracks obtained from 4-bit and 8-bit binned frames during the same time interval as in **b** and **c. e**, Same as **d** but for *y*-tracks. **f**, Full confidence of all beads shifted together for comparison. Bead 1’s confidence is plotted in green, bead 2’s confidence is in red, and bead 3’s confidence is in blue. With this color scheme, white appears where the confidences overlap the most. **g**, The same layout as **f** but for the *y*-positions. **h**, Surface-immobilized fluorescent dye Atto643 with 120 programmed stage movements of ∼170 nm with ∼33 ms dwell times. We use every tenth frame to obtain a pseudo-stroboscopic dataset that allows us to analyze trajectories covering a larger part of the FOV. Shown is the contrast-adjusted sum of all 9,922 40 *µ*s binary frames with the inferred track overlaid. The brightness of Atto643 dyes is about one-tenth the brightness of fluorescent beads, giving less than half the photon arrivals per molecule per 40 *µ*s 1-bit frame as for the 10 *µ*s 1-bit fluorescent beads frames. The background photon counts are also 5 to 10*×* higher. Experimental parameters include: numerical aperture of 1.49, refractive index of 1.52, and emission wavelength of 655 nm. **i**, Full *x*-trajectory confidence with SP^2^T and TrackMate 8-bit tracks overlaid, with zoomed inset; 4-bit TrackMate tracks were excluded due to insufficient brightness. Note the missing segments in the TrackMate track. **j**, Same as **i** for the *y*-track.

Furthermore, since fluorescent beads are known to remain stationary during dwell time, it is possible to sum the binary frames over time (binning) to perform conventional tracking. For this purpose, we binned 1-bit frames to obtain 4-bit and 8-bit frames, which we can analyze using TrackMate(*51*). The tracks generated by TrackMate are also plotted in Figs. 2d and 2e, showing alignment, albeit at a lower spatial resolution, with the SP^2^T results. This strong correlation indicates that SP^2^T achieves a similar spatial resolution while offering a temporal resolution that is two orders of magnitude greater.

Next, we test the robustness of SP^2^T for tracking dim emitters using surface-immobilized Atto643 fluorescent dyes undergoing programmed stage movements. Compared with the fluorescent beads, Atto643 yields approximately one-tenth the number of photons per particle, while the background photon level is 5 to 10*×*higher. We additionally analyze a substantially longer trajectory consisting of 120 programmed 170 nm steps, spanning a greater portion of the FOV than in the bead experiment. The stage was programmed with ∼33 ms dwell times, and we construct a pseudo-stroboscopic dataset by retaining every tenth 40 *µ*s exposure 1-bit frame, resulting in an effectively faster particle movement speed and effective dwell time of only ∼3 ms between successive stage movements. Fig. 2h shows the inferred particle track overlaid on the contrast-adjusted sum of all 9,922 40*×*40 pixel binary frames.

Figs. 2i and 2j show the inferred *x*- and *y*-trajectories overlaid on the confidence in green. We estimate a localization uncertainty of about 90 nm from the confidence, which, while larger than the 16 nm uncertainty obtained for the fluorescent beads, is still smaller than the pixel width of 100 nm and well below the diffraction limit of this system. Also included is an 8-bit TrackMate trajectory obtained by binning sets of 255 1-bit frames. Because of the higher background photon level and lower brightness than in the bead dataset, TrackMate was unable to track from 4-bit binned frames as we did for the beads. Note the lower time resolution and missing segments of the 8-bit TrackMate trajectory, highlighting the limitations of the existing intensity-based paradigm when dealing with low-intensity signals and underscoring the robustness of our approach.

We note that SP^2^T assumes a motion model, which it uses to generate track proposals. The model we select is diffusive motion, which poorly characterizes the directed motion of the stage. Even so, both beads and dyes are tracked with remarkable accuracy. This can be understood by considering that the emission model (*i*.*e*., the probability of the data given proposed tracks) typically informs the likelihood far more than the assumed motion model(*52*). As the particle motion here is not in fact diffusive, inferring a diffusion coefficient is not as meaningful or clearly verifiable; however, we do report the diffusion coefficient inferred alongside the trajectory for the surface-immobilized dye case in Fig. S.6, compared to approximate diffusion coefficient values expected for step-wise trajectories by considering the area covered by the trajectory(*53*). To show that the exact choice of diffusion coefficient prior has little effect on the obtained results, Fig. S.7 shows the tracking and diffusion coefficient results obtained for two different prior parameter choices. We return to diffusion coefficient inference in more detail below, for a diffusing Atto565-labeled ganglioside GM1.

### Short camera exposure for accurate diffusion analysis

Next, we highlight the importance of short camera exposures in accurately estimating diffusion coefficients in single-particle tracking. To demonstrate this, we use the SPAD512^2^ camera to acquire 1-bit image frames of diffusing Atto565-labeled ganglioside GM1 molecules at 50 *µ*s per frame (see Fig. 3a). Gangliosides are glycosphingolipids that self-organize into dynamic, nanoscopic domains within biological membranes. The assembly of nanodomains is driven predominantly by the hydrogen bonding capabilities of their sialic acid residues and the glycan headgroup(*54 –56*), making their mobility a sensitive readout of local membrane organization. To compare across exposure times on identical data, consecutive 1-bit frames are summed to generate higher-bit-depth image stacks with effectively longer exposures. For example, summing 15 frames produces a 4-bit image with an effective exposure of 750 *µ*s, while summing 255 frames yields an 8-bit image with an effective exposure of 12.75 ms (see Fig. 3b). These datasets are then analyzed with SP^2^T, which can handle images of any bit depth, to extract trajectories, estimate diffusion coefficients, and compute jump-distance distributions.

**Figure 3:**
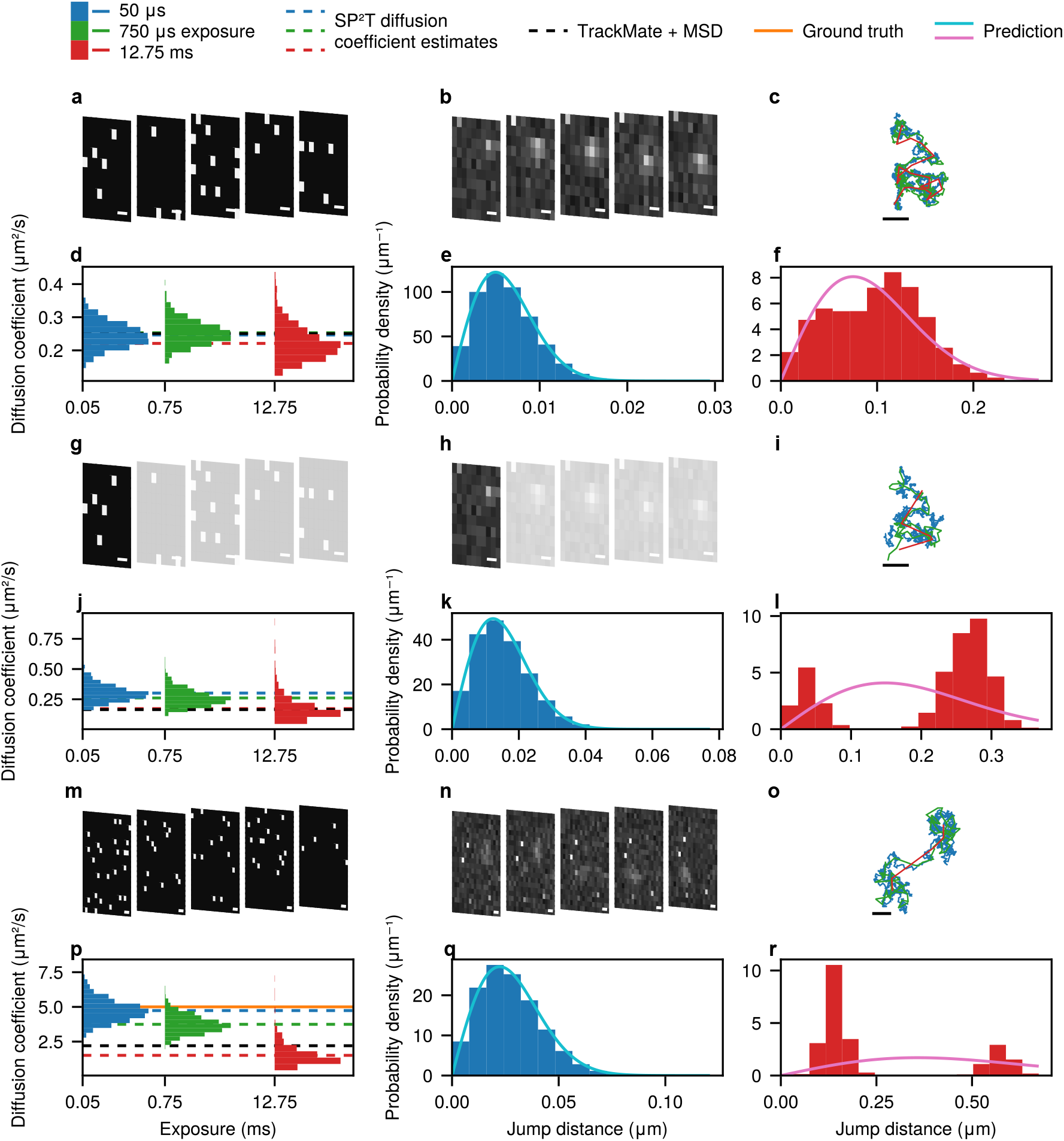
Effects of camera exposure on diffusion coefficient estimates, particle trajectories, and jump-distance distributions. All scale bars in this figure correspond to 200 nm, or the width of two pixels. **a**, Frames of a diffusing Atto565-labeled ganglioside GM1 acquired with the SPAD512^2^ camera (50 *µ*s per frame; see Experimental methods). **b**, Effective 12.75 ms-exposure frames of the same region, generated by summing consecutive short-exposure frames. **c**, Particle trajectories reconstructed by SP^2^T under different exposure conditions. **d**, Estimated diffusion coefficients across effective exposures. SP^2^T results are shown as blue, green, and red dashed lines; TrackMate(*51*) results from 12.75 ms frames are shown as a black dashed line. **e**, Jump-distance distribution between consecutive frames from native 50 *µ*s data compared with the Rayleigh distribution (light blue) predicted from the mean diffusion coefficient estimated by SP^2^T (0.25 *µ*m^2^/s). **f**, Same analysis as in **e**, but using the 12.75 ms frames and the corresponding SP^2^T estimate (0.22 *µ*m^2^/s); Rayleigh prediction shown in pink. **g–h**, Pseudo-stroboscopic dataset generated by retaining only the first frame of every five consecutive frames, reducing the effective frame rate while keeping exposure fixed. **i–l**, Analyses of the pseudo-stroboscopic dataset, arranged as in **c–f. m–r**, Same analyses applied to simulated data with diffusion coefficient 5 *µ*m^2^/s (orange horizontal line in **p**). Simulation parameters match experiments: objective numerical aperture 1.49, pixel size 100 nm, refractive index 1.52, 50 *µ*s exposure, and emission rate of ∼15,000 photons/s (comparable to beads in Fig. 2).

As illustrated in Fig. 3c, shorter exposures (50 *µ*s) produce trajectories with many more steps than those reconstructed from long-exposure frames, reflecting the much higher temporal resolution available when the same total number of 1-bit frames is used. The diffusion coefficient estimates, however, remain consistent across this range (Fig. 3d): the 95% credible interval (CI)—the range containing 95% of the posterior probability mass—spans 0.17 *µ*m^2^/s to 0.32 *µ*m^2^/s at 50 *µ*s and 0.14 *µ*m^2^/s to 0.33 *µ*m^2^/s at 12.75 ms. Both lie about an order of magnitude below the ensemble value of (2.6 *±* 1.3) *µ*m^2^/s reported for this system(*56*). This difference reflects what each measurement averages over: the reported value is an ensemble average across a large field of view, whereas ours derives from a small region of interest containing a single GM1 molecule.

Re-analyzing the same experimental sample with an EMCCD camera (512*×*512 pixels, 50 ms exposure) using TrackPy(*57*) (Fig. S.8) recovers a broad distribution of single-molecule diffusion coefficients, from below 0.01 *µ*m^2^/s to nearly 20 *µ*m^2^/s, with about 18% falling in the 0.1 *µ*m^2^/s to 0.5 *µ*m^2^/s range where our estimate lies. Such breadth is consistent with local inhomogeneities in the supported lipid bilayer (SLB)(*58, 59*).

In contrast to the diffusion coefficient estimates, which remain consistent from 50 *µ*s to 12.75 ms exposures (Fig. 3d), the impact of exposure time is far more pronounced in the jump-distance distribution (displacements between consecutive frames) and its interpretation. As shown in Fig. 3e, the distribution obtained from 50 *µ*s exposures closely matches the Rayleigh distribution predicted from the mean diffusion coefficient estimated by SP^2^T using the same frames. Crucially, this Rayleigh curve is not a fit but a prediction from the inferred diffusion coefficient. At 12.75 ms exposures, the distribution deviates strongly from the same prediction (Fig. 3f). While the diffusion coefficient was itself estimated from these frames under a Brownian model, the failure of that estimator on the same data at longer exposures isolates exposure time as the cause. Such a discrepancy can lead to spurious inference of the diffusive model: the skewed distribution might be misinterpreted as evidence of heterogeneous diffusion or anomalous transport, when in fact it is an artifact of reduced temporal resolution.

To further investigate the impact of limited temporal resolution, we created a pseudo-experimental dataset by retaining only the first frame of every five consecutive frames, mimicking a stroboscopic illumination scheme(*26 –28*) (Figs. 3g to 3i). This reduces the effective frame rate by a factor of five while keeping the exposure per frame constant. Under this setup, the estimated diffusion coefficient falls to a mean of 0.17 *µ*m^2^/s with substantially greater uncertainty (95% CI: 0.07 *µ*m^2^/s to 0.41 *µ*m^2^/s; Fig. 3j), compared with 0.22 *µ*m^2^/s recovered from the full-rate data at the same exposure. Discarding four of every five frames leaves roughly five times fewer trajectory steps, and it is this loss of steps—not the exposure time alone—that drives the underestimate, consistent with recent observations(*24*).

Moreover, the stroboscopic 12.75 ms jump-distance distribution becomes bimodal (Fig. 3l), a feature absent from both 50 *µ*s distributions (Figs. 3e and 3k) and the full-rate 12.75 ms distributions (Fig. 3f). We attribute this newly emergent bimodality to the small number of trajectory steps that survive once frames are discarded. Recovering the correct dynamics under these conditions would require many more trajectories, which reinforces the main conclusion: short exposures yield more reliable diffusion coefficient estimates and jump-distance distributions from the same amount of data.

For comparison, Fig. S.9 presents results obtained when the other four frames in each five-frame block were retained instead of the first, while Fig. S.10 shows results from a different region of interest within the same dataset. Finally, we note that the same form of bias observed in Figs. 3j to 3l can also be reproduced *in silico* with a diffusion coefficient of 5 *µ*m^2^/s, without discarding frames (Figs. 3m to 3r), further emphasizing the need for adequate temporal sampling in single-particle tracking experiments.

These findings raise an important question: How short must the camera exposure be to reliably recover both the correct diffusion coefficient and the underlying diffusive model? Defining a strict threshold is difficult, as it depends on many factors, including the particle’s diffusion coefficient, fluorophore brightness, background noise, camera characteristics, and even the trajectory’s geometry. Nevertheless, our analysis in Fig. 3 reveals a clear trend: shorter exposures consistently produce more accurate and reliable results. Accordingly, we advocate minimizing exposure times wherever possible. This reinforces the importance of moving beyond the localization paradigm, which inherently relies on finite exposures to improve per-frame fits, and underscores the dual need for ultrafast imaging hardware such as SPAD arrays, together with tracking algorithms that can extract precise information from low-bit-depth, high-temporal-resolution data.

### Multiplexing

Having demonstrated SP^2^T’s tracking of real diffusing molecules at microsecond temporal resolution, we next evaluate its ability to perform multiplexed tracking. We consider three complementary challenges: tracking multiple emitters across a large FOV, simultaneously tracking 100 emitters at 100 *µ*s exposure, and distinguishing emitters with overlapping PSFs. Together, these tests evaluate SP^2^T’s ability to track not only isolated particles from single-photon frames, but also the multi-emitter conditions relevant to biological imaging.

We first demonstrate multiplexed tracking experimentally by analyzing Cy3B-labeled pro-aerolysin monomers diffusing on an SLB across the full 512*×*512 pixel FOV, captured using the SPAD512^2^ camera from Pi Imaging. Aerolysin is a protein involved in pathogenic infections caused by *Aeromonas* species(*60*) and widely used in biosensing applications(*61*). The mechanism of membrane association and pore formation has never been captured dynamically. Here, most molecules appear immobile, suggesting strong interactions with the membrane, while a small fraction diffuses faster, indicating weaker association. This dataset comprises 1,000 frames, with each molecule emitting, on average, approximately two photons per frame. Frames were captured with an exposure time of 100 *µ*s, allowing us to evaluate SP^2^T’s performance under realistic, high-throughput imaging conditions.

Figure 4a presents the sum of all 1,000 frames for illustrative purposes only. Before processing this dataset, we first selected 5,000 frames (the 1,000 frames used for analysis, along with the subsequent 4,000 frames) and computed a temporal average to serve as a background calibration. This approach ensures that stationary particles are treated as part of the background and thus excluded from tracking, allowing SP^2^T to focus on dynamic molecules. We ascribe the stationarity of the bright spots (molecules) in this dataset to incomplete activation of the aerolysin molecules—specifically, the failure to cleave their Cterminal peptides(*62, 63*). Using this background calibration, SP^2^T identified and tracked seven diffusing aerolysin molecules.

**Figure 4:**
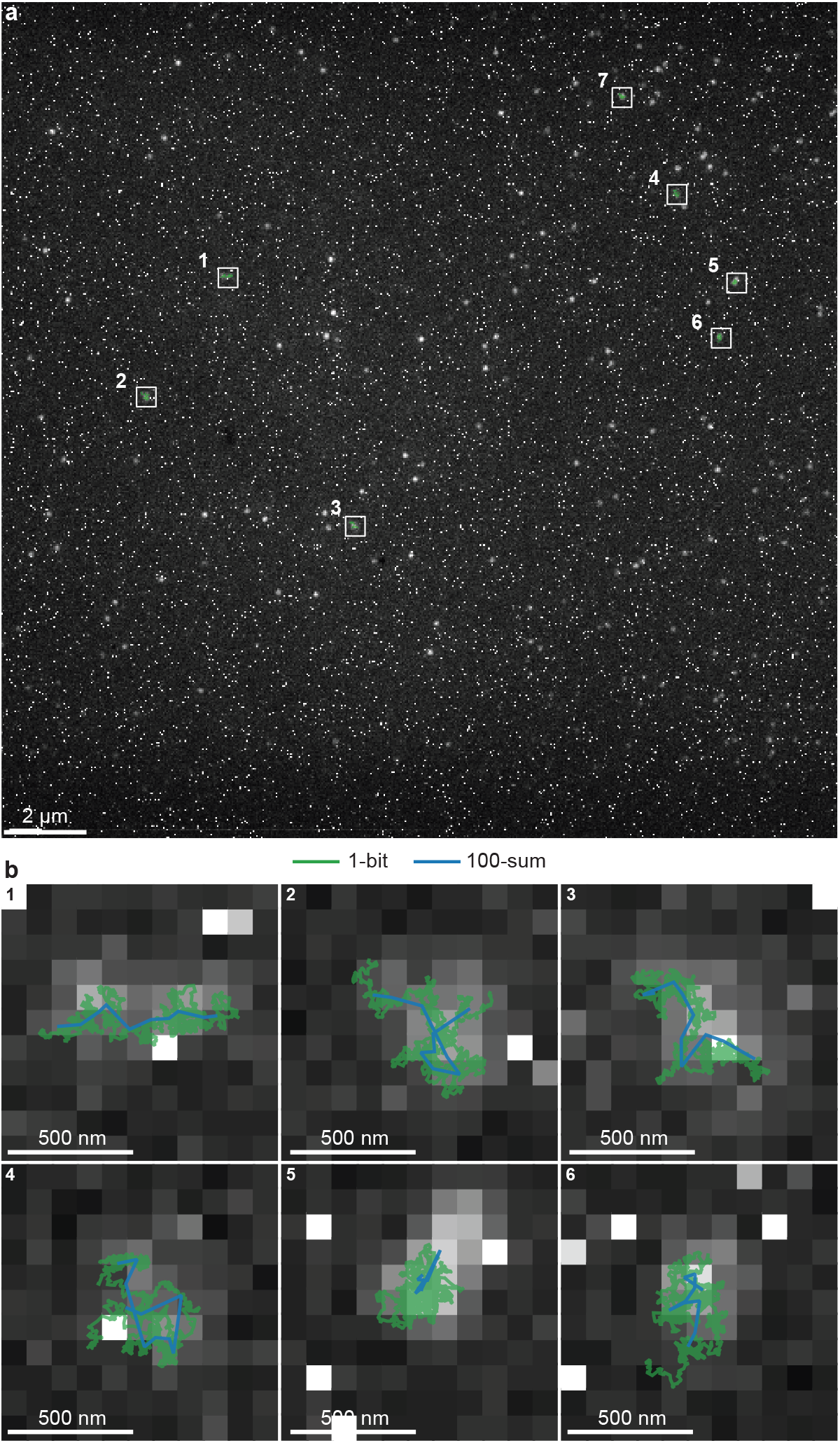
Tracking diffusing aerolysin molecules across a 512*×*512 pixel FOV of the SPAD512^2^ camera from Pi Imaging. The dataset consists of 1,000 frames with 100 *µ*s exposures. **a**, The contrast-adjusted sum of all binary frames for the full FOV is shown for illustrative purposes. Seven aerolysin molecules identified as nonstationary are highlighted with white boxes. **b**, Zoomed-in views of the selected regions containing six of these seven molecules, for optimal figure clarity only. SP^2^T tracks are overlaid in green, while particle tracks extracted using TrackMate from 100-sum frames (10 frames of 100 consecutive binned frames) are shown in blue. Experimental parameters include: numerical aperture of 1.49, refractive index of 1.518, and emission wavelength of 571 nm.

In Fig. 4b, we show zoomed-in views of six of the seven tracked particles (for clarity), comparing the 1-bit particle tracks inferred by SP^2^T with those extracted using TrackMate from 100-sum frames, where each frame is generated by summing 100 consecutive frames (10 binned frames in total). Consistent with our previous results, SP^2^T achieves particle tracking with a 100-fold improvement in temporal resolution over TrackMate while maintaining a spatial resolution comparable to that achieved with TrackMate on 100-sum frames. Fig. S.11 contains a more in-depth analysis of a single diffusing aerolysin molecule dataset with the same experimental conditions.

Having demonstrated multiplexed tracking experimentally, we next use synthetic data with known ground truth to test SP^2^T at substantially larger particle numbers. We simultaneously track 100 diffusing particles from 1,000 1-bit, 128 *×* 128 pixel frames acquired at 100 *µ*s exposure. Simulation parameters, including emitter brightness, diffusion coefficient, and background photon levels, are chosen to mimic the aerolysin dataset in Fig. 4. Fig. 5a shows the binned sum of all frames for the full FOV for illustrative purposes, along with the ground truth and inferred trajectories for all 100 particles in orange and green, respectively. Fig. 5b zooms into a single particle, showing the consistency of the ground truth and inferred trajectory. Fig. 5c shows a single 1-bit frame from the dataset for the same zoomed-in region, for the particle location indicated by the orange cross, illustrating the need to consider photon arrivals from the whole dataset to track.

**Figure 5:**
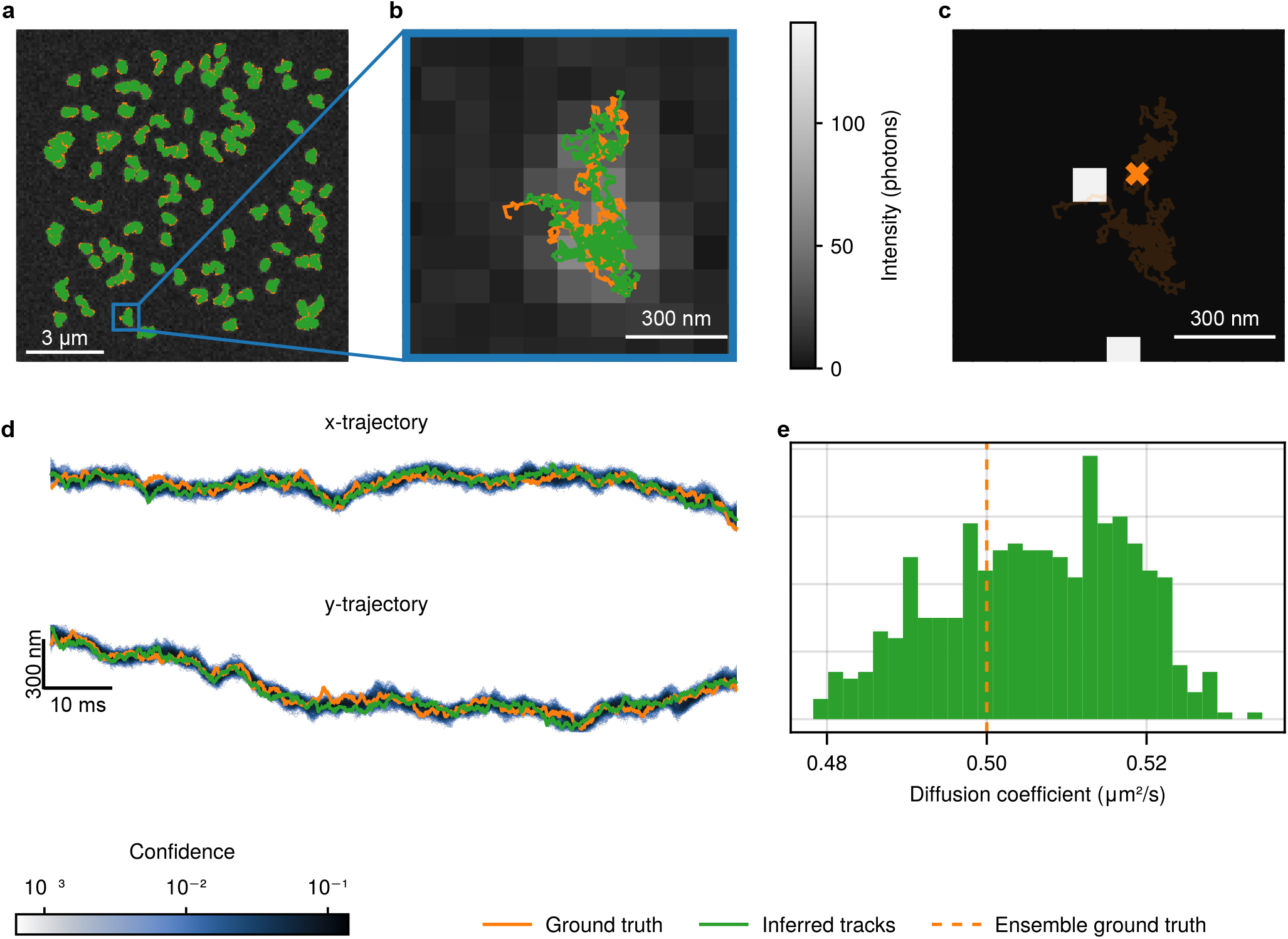
Multiplexed tracking of 100 particles from 1,000 100 *µ*s 1-bit frames. **a**, Binned 128*×*128 frames of the simulation of 100 particles, for illustrative purposes, using emitter brightness (10,000 photons per second per emitter) and background photon rate (14,000 photons per *µ*m^2^ per second) to match the aerolysin data in Fig. 4, with ground truth and an inferred trajectory overlaid. **b**, Zoom-in to a single particle, with the inferred trajectory in green and the ground truth in orange, with a scale bar of length 300 nm, or 3 pixels. **c**, For the same region, a single 1-bit frame showing two photon arrivals for the point in the trajectory indicated by the orange cross. **d**, Overlap of the ground truth trajectory and credible interval indicates excellent match between inferred and ground truth, with darker indicating greater confidence, for both the *x*- and *y*-trajectories. **e**, The posterior distribution of inferred diffusion coefficient includes the true ensemble diffusion coefficient of 0.5 *µ*m^2^/s. Other simulation parameters: numerical aperture of 1.49, refractive index of 1.52, emission wavelength of 665 nm, and pixel size of 100 nm.

For the particle shown in Fig. 5b, Fig. 5d shows the inferred and ground truth *x*- and *y*- trajectories overlaid on the confidence, calculated from posterior samples. Empirical coverage is close to nominal, with the ground truth trajectory falling within the *x*- and *y*-trajectory 95% credible intervals 92% and 94% of the time, respectively. Particle tracks are inferred with a mean localization error of 48 nm, consistent with our previous results, despite the far larger number of particles present. We highlight that tracking so many particles with sub-pixel accuracy across a large FOV at single-photon arrival temporal resolution extends single-photon tracking to the crowded, large FOV regime relevant to biological samples. Additionally, Fig. 5e also shows the posterior distribution of the learned diffusion coefficient for the ensemble of particles, which is consistent with the ensemble ground truth value of 0.5 *µ*m^2^/s. Fig. S.12 shows tracking of 50 diffusing particles at 50 *µ*s exposure time, and Fig. S.13 and Fig. S.14 show the convergence of the 100 and 50 particle cases.

Finally, we test multiplexing in one of the most challenging regimes for particle tracking, relevant to dense biological samples: two emitters with overlapping PSFs. As the emitters approach one another, the photons collected across the detector become increasingly consistent with those produced by a single brighter emitter. Conventional localization methods cannot break this degeneracy from a single frame, potentially merging the emitters into one localization or producing mislinked trajectories. SP^2^T, in contrast, jointly evaluates candidate emitter numbers and trajectories over the entire 1-bit image stack, leveraging the temporal information contained in photon detections across frames, together with the expected photon counts from its calibrated per-emitter brightness model. Thus, in this framework, the total photon budget itself becomes informative in distinguishing between candidate emitter configurations. We therefore test whether SP^2^T continues to infer two distinct emitters as their separation is steadily reduced, including the limiting case of complete spatial overlap. To perform this test with known ground truth, we generate synthetic 1-bit data using two bead trajectories inferred in Fig. 2, simulated with the detector’s measured per-pixel dark counts and emitter brightness matched to the aerolysin data in Fig. 4 (see Methods). We systematically vary their relative spatial offset to generate datasets with decreasing emitter separation, from 400 nm down to 0 nm in 100 nm steps. Zero separation represents the maximally degenerate case and is more challenging than two trajectories that only transiently cross: because the two bead trajectories arise from the same programmed stage motion, the emitters remain nearly coincident throughout the entire dataset. Consequently, the simulated photon detections are consistent with a single emitter of approximately twice the brightness, leaving the calibrated per-emitter brightness as the key constraint distinguishing the two hypotheses.

Figs. 6a, 6e, 6i, 6m and 6q show the summed 1-bit frames at each separation. Two distinct intensity maxima are visible at 400 nm (Fig. 6a); by 300 nm (Fig. 6e) they have begun to merge, and at 200 nm and below (Figs. 6i, 6m and 6q) the summed image is a single diffraction-limited spot in which no second emitter is apparent by eye. Conventional analysis of these frames would report one particle. SP^2^T, by contrast, recovers two separate trajectories at every separation, including complete overlap (Figs. 6q to 6t). Across all five conditions, the MAP tracks follow the ground truth to within 50 nm (half a pixel) on average. One might expect precision to degrade as the emitters converge, since a photon detected at a given pixel becomes progressively less diagnostic of which emitter produced it. In practice we see no such degradation: the marginal posterior width—our measure of localization uncertainty—is essentially unchanged between 400 nm separation and complete overlap.

**Figure 6:**
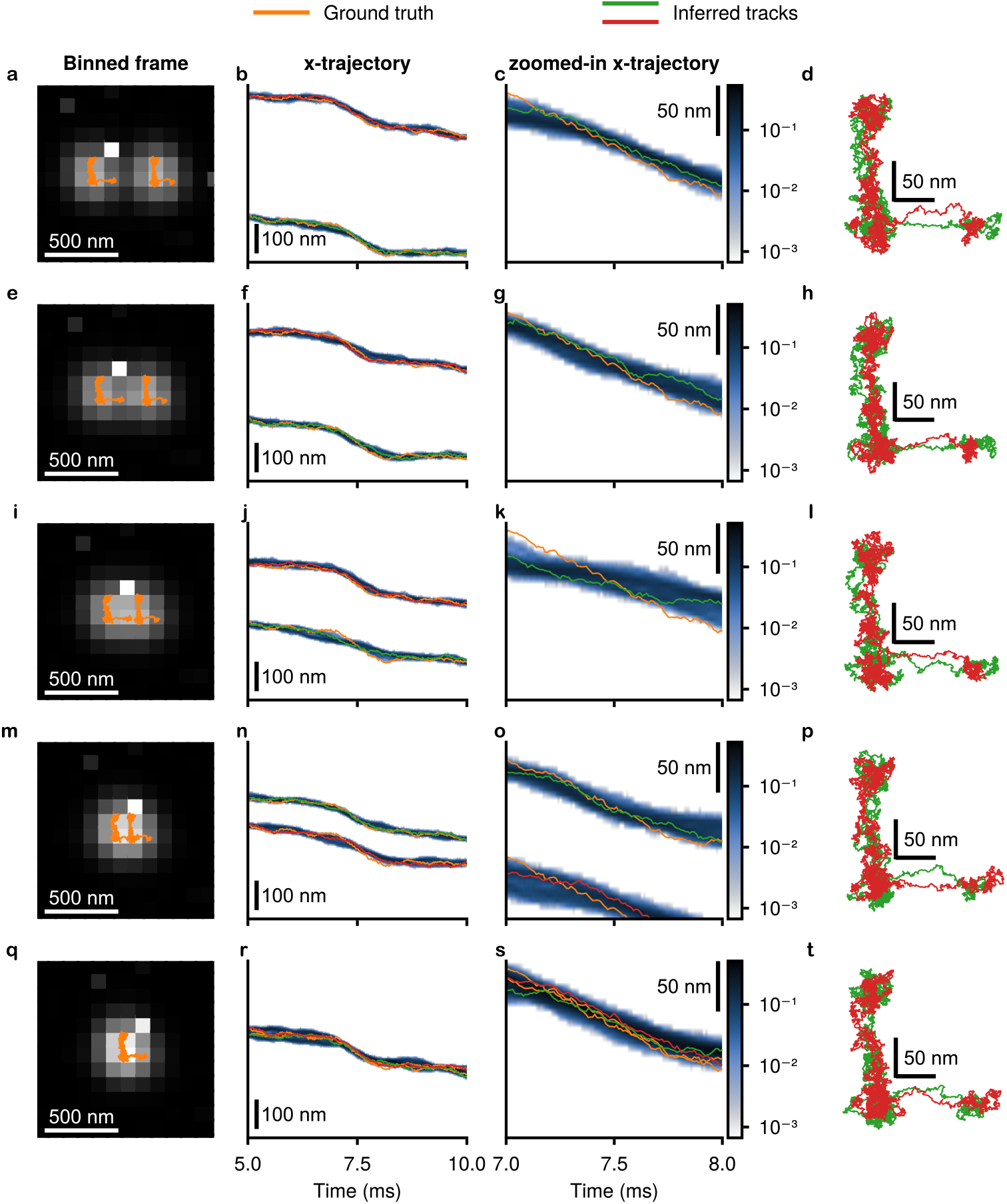
Resolving two emitters as their trajectories are brought together. Two bead trajectories inferred in Fig. 2 were used as ground truth and placed side by side at a series of decreasing mean separations. 1-bit frames were then simulated from the model in Methods using a numerical aperture of 1.45, refractive index 1.515, emission wavelength 665 nm, 10 *µ*s exposure, 5,750 frames, per-pixel dark counts measured from the detector used in Fig. 2 with 100 nm pixels, and emitter brightness matched to Fig. 4. Inference was run nonparametrically, with the number of emitters inferred. Rows correspond to mean separations of 400 nm (**a–d**), 300 nm (**e–h**), 200 nm (**i–l**), 100 nm (**m–p**), and complete overlap, 0 nm (**q–t**). **a**, Sum of all 1-bit frames, shown for illustrative purposes, with the two ground-truth trajectories overlaid in orange; scale bar 500 nm. **b**, The two inferred *x*-trajectories between 5 ms and 10 ms; ground truth in orange, MAP tracks in green and red. Shading gives the marginal posterior probability that the emitter lies within a given 10 nm bin, on a logarithmic color scale; scale bar 100 nm. **c**, The same tracks between 7 ms and 8 ms, zoomed in both time and position. Because the displayed position range is held fixed across rows, only one of the two trajectory bands falls within it at the larger separations; scale bar 50 nm. **d**, The two inferred tracks, with ground truth omitted, after a constant translational shift, so that their consistency with one another can be assessed; scale bar 50 nm.

With the emitter brightness calibrated, the posterior over emitter number places all of its mass on two emitters at all tested separations, including complete spatial overlap. Importantly, in the fully overlapping case, SP^2^T is not spatially resolving two distinct spots— the summed image contains only one. Rather, the total photon budget, interpreted through the calibrated per-emitter brightness model, provides the information needed to favor the two-emitter hypothesis over a single brighter emitter. These results demonstrate that SP^2^T can maintain multiplexed tracking even as emitters become spatially crowded and their PSFs strongly overlap. This extends single-photon tracking toward the dense multi-emitter conditions encountered in biological imaging.

## Discussion

We have shown how to track emitters directly from binary single-photon detector frames at microsecond exposures. By jointly analyzing 1-bit image stacks and circumventing the localization-then-linking paradigm, SP^2^T pushes temporal resolution from milliseconds to microseconds while maintaining sub-pixel localization precision. It recovers the correct particle number and accurate trajectories across diverse datasets: from fluorescent beads and dyes on a pre-programmed stage, through Cy3B-labeled aerolysin monomers on SLBs and Atto565-labeled ganglioside GM1 molecules, to 100 particles tracked at 100 *µ*s temporal resolution.

We compared SP^2^T against TrackMate(*51*), a widely used single-particle tracking tool, on binned versions of every dataset here. TrackMate and related tools(*64, 65*) achieve high spatial resolution from long-exposure frames, but remain fundamentally committed to localization itself. SP^2^T provides tracks of comparable localization precision while circumventing that paradigm and achieving one to two orders of magnitude higher temporal resolution, eliminating blurring artifacts and spanning both slowly diffusing organelles(*49*) and faster-diffusing drug nanoparticles (Fig. 1b). This opens faster processes, such as vesicle transport and nanoparticle delivery, to study at previously inaccessible temporal resolution. Multiplexing further allows many particles to be tracked across a large FOV, including closely spaced emitters, bringing crowded biological samples within reach.

Beyond the choice of tracking algorithm, our results highlight a more fundamental issue: long camera exposures systematically bias estimates of diffusion parameters and of the underlying diffusive model (Fig. 3), especially for fast-moving particles, regardless of the tracking method used. We therefore advocate the shortest practicable exposures, analyzed with SP^2^T, which handles low-bit-depth data without frame summation. This hardware-software pairing stays accurate even in photon-starved conditions, substantially extending the dynamic range accessible to single-molecule studies.

Although SP^2^T offers clear advantages, it has limitations, both computational and methodological. Computationally, the challenge is frame count: operating directly on binary frames, SP^2^T may face up to 255 times more frames than an 8-bit equivalent, and analyzing them jointly becomes demanding for large datasets. We address this with GPU parallelization and flexible parametric inference.

First, the likelihood calculation, the most intensive component, is parallelized in a two-layer structure and executed on a GPU (see Methods), giving up to a 50-fold speedup on an NVIDIA GTX 1060 over an Intel i7-7700K CPU.

Second, SP^2^T runs in either nonparametric mode, jointly inferring particle number and trajectories, or parametric mode, where the user supplies the particle count and bypasses that inference. Analyzing nonparametrically first and then rerunning parametrically with the inferred count therefore gives a several-fold gain, about five-fold for ten emitters on the hardware above, though the exact figure depends on emitter number. Initializing SP^2^T with tracks from long-exposure frames yields a further twofold speedup. Users can thus balance efficiency against data complexity.

Together, these optimizations allow SP^2^T to process more than 5,000 binary frames of 50*×*50 pixels in a few hours on a GTX 1060. Because the likelihood is parallelized across both frames and particles, cost stays approximately constant as either grows, until the GPU cores saturate.

Two further limitations are methodological. SP^2^T currently takes the photophysical states 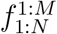 as a fixed input rather than inferring them, so blinking and bleaching must be specified in advance. Separately, distinguishing emitters well below the diffraction limit rests on the calibrated per-emitter brightness (Fig. 6); in practice, brightness varies with dipole orientation, axial position, and photophysical state, so the accuracy of that calibration bounds how far this capability extends. Both point toward the same extension: bringing quantities SP^2^T now treats as fixed inputs into the inference itself.

Several extensions follow naturally from the framework. SPDs are increasingly replacing integrative detectors across applications including fluorescence lifetime microscopy(*32, 36*), image-scanning microscopy(*33, 35*), and Förster resonance energy transfer(*36*), and because SP^2^T operates directly on single-photon data it can absorb the additional information they provide. For detectors that record photon arrival times, the observation model could extend beyond the binary measurements of equation (5) to arrival-time distributions. The LIN-Cam(*66*) and TCSPC-capable SPAD arrays(*35*), with pulsed excitation, already capture the lifetimes needed to separate fluorophores by decay profile(*67*), enabling simultaneous tracking and species identification in complex environments. Expanding the latent-state model to infer photophysical states jointly with trajectories would remove the need to specify them in advance, while modeling brightness as a function of dipole orientation, axial position, and photophysical state would sharpen the calibration on which emitter separation depends. Specialized 3D PSFs, obtained by modifying the integrand in equation (4), would improve three-dimensional tracking. The framework also adapts to non-array SPDs such as the NCam(*68–70*), which remove the need for spatial binning just as we avoid temporal binning here, and could enable truly bin-free imaging in both space and time.

More broadly, SPAD arrays and related architectures that resolve low photon counts per pixel enable new experimental designs, but they also demand that we rethink localization: single-photon data cannot be analyzed with frame-based paradigms built on high photon counts and smooth per-pixel intensities. SP^2^T meets this need by tracking directly from photon arrival events. Both these detectors and the computational methods that exploit them remain at an early stage, and continued advances in hardware, algorithms, and software should extend their reach to dynamic biological events at previously inaccessible spatiotemporal scales.

## Methods

### Theoretical framework

This section outlines the mathematical framework of SP^2^T. For notational simplicity, we first define the following terms: *N* represents the number of frames, *P* denotes the number of pixels per frame, 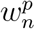 refers to the data collected at pixel *p* on frame *n, M* indicates the number of particles, 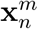 represents the *m*-th particle’s position at frame *n*, and MSD is the mean squared displacement, as later defined in equation (10). With these definitions, we can express all collected data as 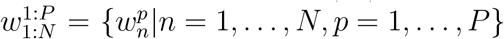. Similarly, the collection of all particle tracks is denoted as 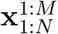.

As discussed in the sections above, the goal of SP^2^T is to determine all variables of interest (including the number of light-emitting particles, particle tracks, and the MSD) along with the corresponding credible intervals. We achieve this by constructing the full probability distribution, known as the posterior, 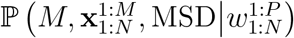, representing the probability distribution of all variables of interest given the collected data. Following Bayes’ theorem, this posterior can be expressed as

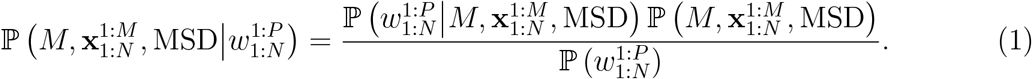

The first term in the numerator, 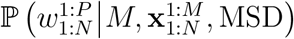, represents the likelihood reflecting the probability distribution of the data given the variables of interest. This term usually describes the system’s physics (the model for photon detection in this context) and realistic complications such as hot pixels. The second term, 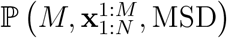, known as the prior, and ensuring the MSDremains non-negative. The denominator, 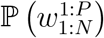, is referred to as imposes additional constraints on the variables of interest, such as a particle’s motion model the evidence and is generally treated as a normalization constant, as it does not depend on any variable of interest.

In the following sections, we will establish the base likelihood using only the photon detection model. Next, we will generalize this base likelihood to account for the aforementioned realistic complications and then discuss the prior we place on the variables of interest. Finally, we will briefly introduce our approach to constructing and sampling the posterior.

### Base likelihood

As introduced above, the likelihood 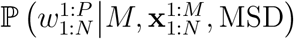 represents the probability distribution of the data given the number of particles, the particle tracks, and the MSD. To express this likelihood mathematically, we first observe that once all particle tracks are known, neither the number of particles nor the MSD affects photon detection. This observation allows us to simplify 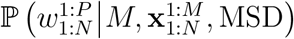 to 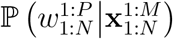.

Furthermore, considering that we are dealing with fluorescently emitting particles whose lifetimes (on the order of nanoseconds(*71*)) are significantly shorter than the frame period (at least 10 *µ*s), the probability that an emitting particle excited during the exposure period of one frame will result in a photon detection in the next frame is negligible (less than 10^−200^, assuming a 10 ns lifetime and excitation occurring in the middle of an exposure). Therefore, it is reasonable to assume that particle positions at frame *n* do not influence other frames. This assumption allows us to further simplify the likelihood as a product

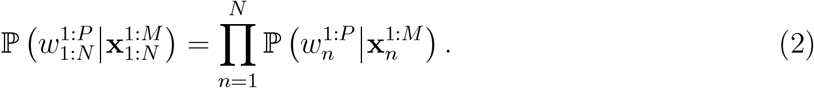

Similarly, since the pixels of a detector usually operate independently, we can express the per-frame likelihood 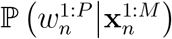 as a product over the individual pixel likelihoods

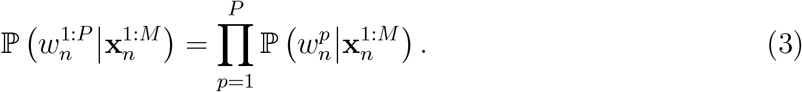

The per-pixel likelihood 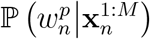 is influenced by four key factors: the PSF of the optical system, the pixel area *A*, the brightness of the particles *h*, and the number of particles *M*. The PSF characterizes how light from a point source is distributed across the detector(*72*). Brightness, in turn, is the expected number of photons arriving at a pixel perfectly aligned with the PSF of a particle located at the focal plane. The total number of photons received by pixel *p* from all emitting particles, denoted as 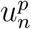, can be expressed as

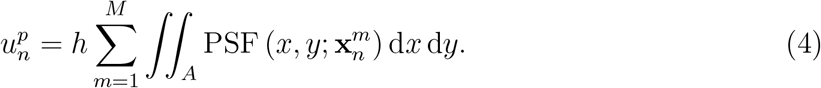

Since a single-photon detector can only distinguish between the absence of photons (false) and the presence of at least one photon (true), the measurement 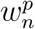 must be sampled from a Bernoulli distribution. The parameter of this Bernoulli distribution is determined by the probability of detecting at least one photon, which is given by 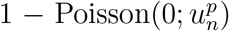. Here, 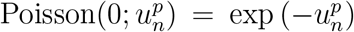 represents the probability of detecting zero photons. Consequently, the probability of detecting at least one photon is 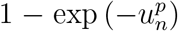, and the corresponding mathematical expression is

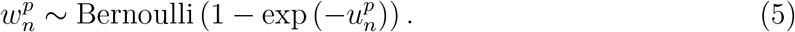

Thus, the overall likelihood can be obtained by combining equations (2) to (5).

### Photophysical states

A crucial factor to consider is the photophysical state of a particle: whether it is in a bright state, actively emitting photons, or in a dark state, where no emission occurs. We define the photophysical state of particle *m* at frame *n* as a binary variable 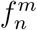, where 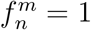 indicates an emitting particle and 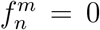 represents a non-emitting state. Currently, SP^2^T takes 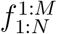 as an input and treats these states as fixed throughout the analysis. Incorporating 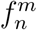 into equation (4) modifies the expected intensity 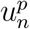 at pixel *p* in frame *n* as follows

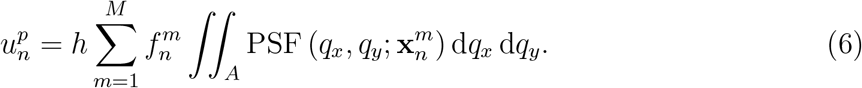

Here, *q*_*x*_ and *q*_*y*_ are dummy variables used to perform the integration over the pixel area.

### Dark counts

Single-photon detectors are known to exhibit dark counts(*35*), which are false detections of photons that occur even in the absence of light. These dark counts, specific to each pixel, arise due to thermal noise and other intrinsic factors within the detector. As a result, the likelihood model must be adjusted to account for these dark counts, which contribute to an additional noise source and affect the accuracy of photon detection. To do so, we introduce a quantity that represents the expected number of dark counts per frame exposure, DC^*p*^, and we modify equation (6) as follows

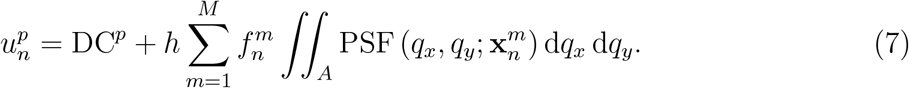

### Other pixel defects

Similar to dark counts, pixels of a single-photon detector can also carry other defects. For example, some pixels may be imperfectly connected to the circuit and, therefore, not produce any signal during an experiment. We therefore allow users to set a Boolean label, *F*^*p*^, which is zero when the corresponding pixel has a defect and one otherwise, to determine whether a pixel should be excluded from the likelihood calculation. Specifically, we change equation (3) to

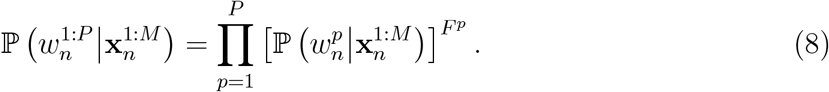

### Full likelihood

To summarize the sections above, we present the mathematical form of the full likelihood here:

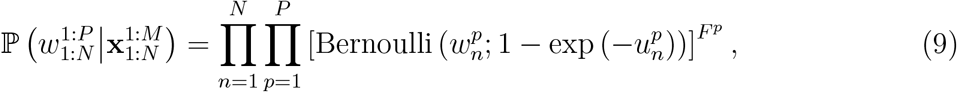

where the expected photon count *u*^*p*^ is given by equation (7).

### Motion model prior

Now that we have established the full likelihood 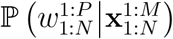, we briefly introduce the motion model prior so that the full posterior can be constructed. As mentioned earlier, we characterize the motion of a particle using the MSD. Specifically, we choose

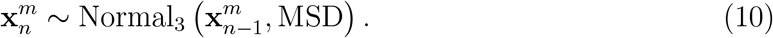

### Posterior

According to equation (1), we can now write the full posterior as

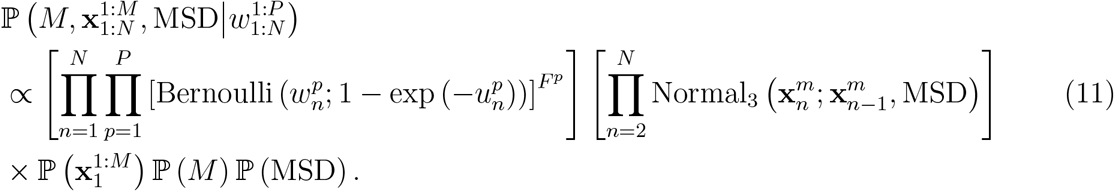

Here, the last three priors 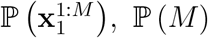, and P (MSD) are chosen for computational convenience. As they primarily define the support of each variable and do not otherwise encode critical physics, we introduce them in the Supplementary Information.

### Parallelization

We parallelize the likelihood calculation at two levels: across frames, and across pixels within each frame. Conceptually, this parallelization scheme stems from the notion that the full likelihood can be expressed as the product of per-pixel likelihoods across all pixels and frames. As an important technical point, these per-pixel likelihoods are independent when conditioned on particle tracks 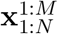 and per-pixel boolean label *F*^*p*^ as shown in equations (8) and (9) so long as SPD pixels are uncorrelated. This independence allows us to efficiently parallelize calculations. Inspired by parallelization schemes across Bayesian applications(*73–76*), we specifically tailor our framework to SPDs to achieve the best possible computational efficiency.

### Experimental methods

#### SPAD array

Experiments were performed on a custom-built widefield single-molecule fluorescence microscope equipped with a large, time-gated 512*×*512 SPAD array (SPAD512^2^, Pi Imaging), in which each pixel can detect single photons, described previously(*77*). This commercial version of the SwissSPAD^2^ camera was developed by the AQUA group (of Prof. Charbon) at EPFL. The camera captures binary images at an exceptional frame rate of up to 97,700 frames per second. A supercontinuum pulsed laser source was used for illumination, capable of emitting arbitrary wavelengths in the visible range. We used either epi-illumination or total internal reflection illumination, allowing monitoring thousands of individual molecules simultaneously within an ≈50*×*50 *µ*m^2^ FOV. The detection path can be swapped for instrument comparison with a conventional EMCCD camera.

### Beads and dye on a moving stage

Widefield imaging for the datasets of fluorescent beads and dyes fixed to a moving stage was performed on a second custom-built fluorescence microscope, as described in Ronceray *et al*.(*78*). Briefly, we imaged surface-immobilized fluorescent beads while following a programmed trajectory set by the motion of the sample placed on a piezoelectric stage (Mad City Labs). Atto643 fluorescent dyes were imaged under the same programmed-trajectory scheme using a motorized stage (Tango Desktop, Märzhäuser).

### Materials

We purchased 1,2-dioleoyl-sn-glycero-3-phosphocholine (18:1Δ9-CisPC, DOPC) dissolved in chloroform, Cholesterol and Brain SM Sphingomyelin from Avanti Polar Lipids, HEPES from Chemie Brunschwig AG, glucose and GM1 ganglioside from Sigma-Aldrich,and Atto643maleimide (AD-643-41, ATTO-TEC/Leica Microsystems) used without further conjugation.

### Aerolysin sample preparation

Full-length wild-type aerolysin protein in the pET22b vector with a C-terminal hexa-histidine tag was used. Cysteines were introduced into specific pore regions by mutating amino acids such as G68, enabling labeling and tracking conformational changes during pore formation(*63*). The mutants were generated by Genscript using the wild-type plasmid as a template, as was previously done for the production of 26 other aerolysin mutants(*79*). Here we followed protocols for aerolysin production, purification, labeling, and preparation of supported lipid bilayers already established in the Radenovic lab(*77, 79*).

### Ganglioside sample preparation

For these measurements, we focused on the diffusing GM1 (monosialotetrahexosylganglioside, a glycosphingolipid with a single sialic acid residue known for its essential role in membrane organization and cell signaling) within DOPC-supported lipid bilayers. Supported lipid bilayers were prepared using 1,2-dioleoyl-sn-glycero-3-phosphocholine (DOPC) / cholesterol / sphingomyelin / GM1 (61:25:10:4, wt%) with 0.5% Allyl-PEG5000, and GM1–Atto565 was incorporated at a lipid-to-dye ratio of 1:15,000,000 to optimize imaging conditions. GM1 was a kind gift from Dr. Ilya Mikhaylov, the synthesis of which is described in Svobodová *et al*.(*56*). Lipid films were generated by evaporating chloroform-dissolved components under vacuum, rehydrated in buffer (10 mM HEPES, 10 mM NaCl, 170 mM glucose, pH 7.4, 1 mM total lipid), and vortexed thoroughly. SUVs were obtained by 90 min sonication and separated by centrifugation at 16,500*×*g for 20 min. To form supported bilayers, 60 *µ*L of SUVs, 140 *µ*L buffer, and 0.6 *µ*L 1 M CaCl_2_ were incubated in coverslip chambers for 90 min, followed by extensive washing (20*×*100 *µ*L buffer) to remove excess material.

## Declarations

### Availability of data and materials

The datasets analyzed during the current study are available from the corresponding author on reasonable request. Our code is publicly available at https://github.com/LabPresse/SP2T.jl

### Authors’ contributions

S.P., L.W.Q.X. and A.R. conceived the study. N.R., M.F.M., T.B., D.S., R.S., M.H. and A.R. performed the experiments and collected the data. L.W.Q.X. and J.C.S. developed the software and carried out the analysis. L.W.Q.X., S.P., A.R., M.F.M. and N.R. wrote the original draft. All authors read and approved the final manuscript.

## Acknowledgments

The authors thank Vytautas Navikas for his effort in fine-tuning the figures in this manuscript. L.W.Q.X. thanks Zachary H. Hendrix for helpful literature suggestions throughout.

