## Supplementary Information for "Tracking Without Linking or Localization: Single-Photon Tracking"

#### A Additional figures

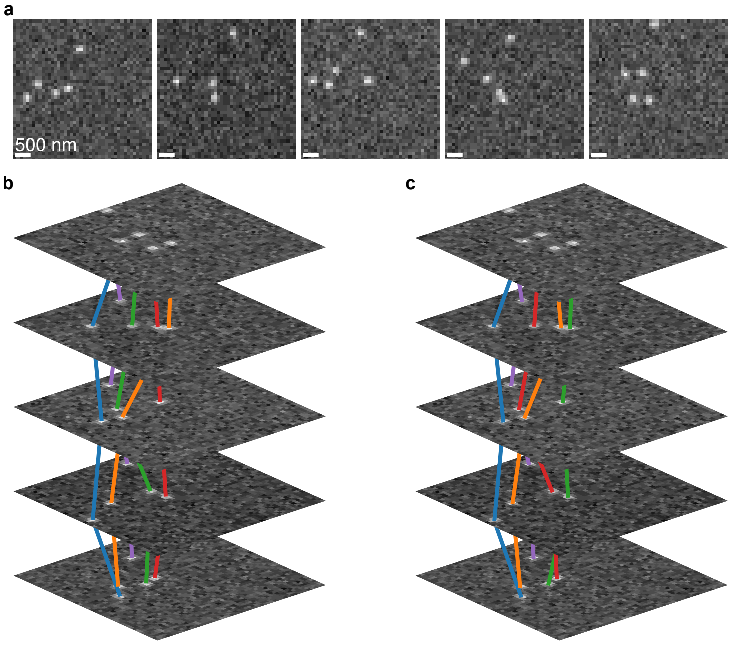


Figure S.1: Stroboscopic illumination can lead to mislinking in particle tracking for fast-diffusing molecules. **a**, Simulated dataset consisting of five consecutive frames, each with a 4 ms illumination window and a 40 ms frame period. Five particles diffuse at $2\,\mu m^2/s$, with imaging parameters: numerical aperture 1.49, refractive index 1.52, emission wavelength 571 nm, and pixel size 100 nm. **b**, Ground truth trajectories, color-coded by particle identity. **c**, Tracks reconstructed using TrackMate(*1*). Mislinking occurs in two of the five frames: in frame 2, the red and green tracks are swapped; by frame 4, the red, green, and orange tracks are permuted. These errors result from long dark intervals between illumination pulses, during which fast-moving particles displace significantly, complicating accurate linking.

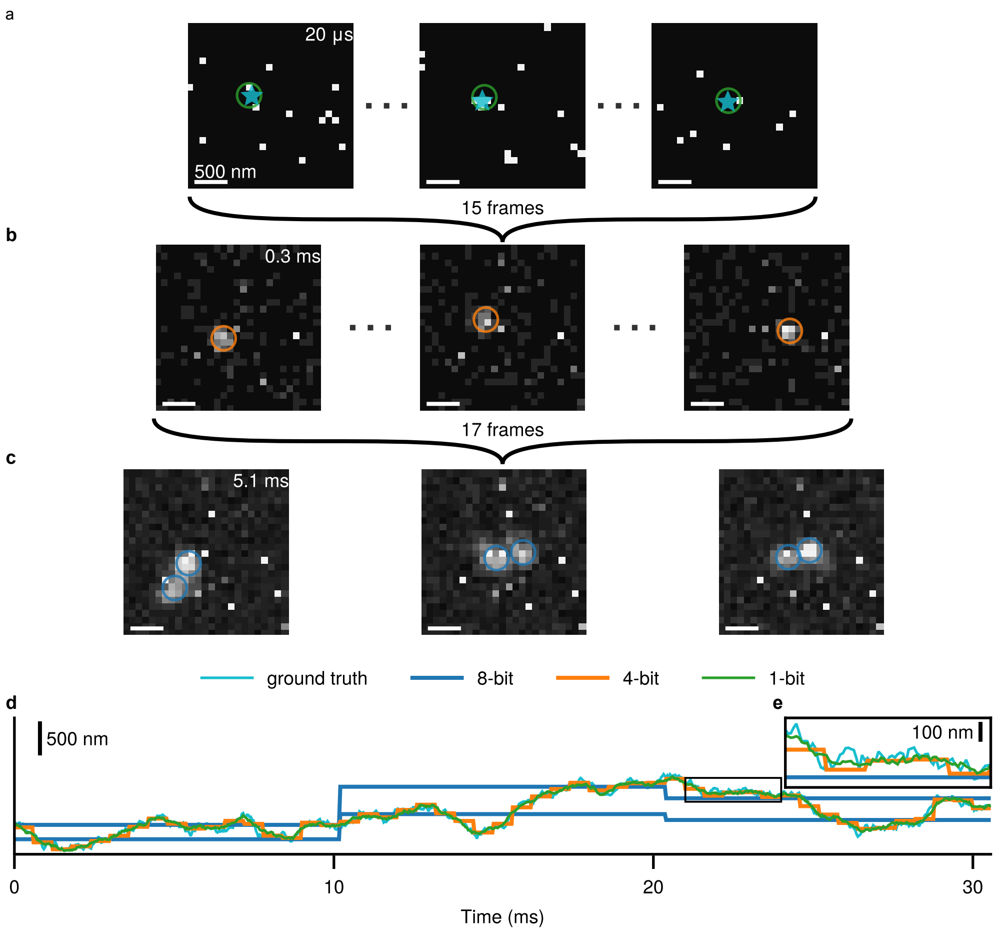


Figure S.2: **a**, Example 1-bit frames with molecule positions (with ground truth shown as a star) extracted by our proposed framework here, indicated by green circles. 15 1-bit frames are summed to produce a 4-bit frame. **b**, Example 4-bit frames with target positions localized using TrackMate(*1*), indicated by orange circles. Seventeen 4-bit frames are summed to create an 8-bit frame. **c**, Example 8-bit frames, where blue circles show particle positions detected by TrackMate. Due to severe blurring artifacts, TrackMate incorrectly identifies two instead of one target. For visualization purposes, the contrast in frames of lower bit-depth is enhanced in **a**–**c**. **d**, Comparison of tracks obtained from frames with different bit-depths: 1-bit tracks are extracted using the proposed framework, while 4-bit and 8-bit tracks are obtained using TrackMate. **e**, Zoom in onto the boxed region in **d**. Simulation parameters include: diffusion coefficient of $10\,\mu\text{m}^2/\text{s}$, numerical aperture of 1.49, refractive index of 1.52, emission wavelength of 571 nm, and pixel size of 100 nm.

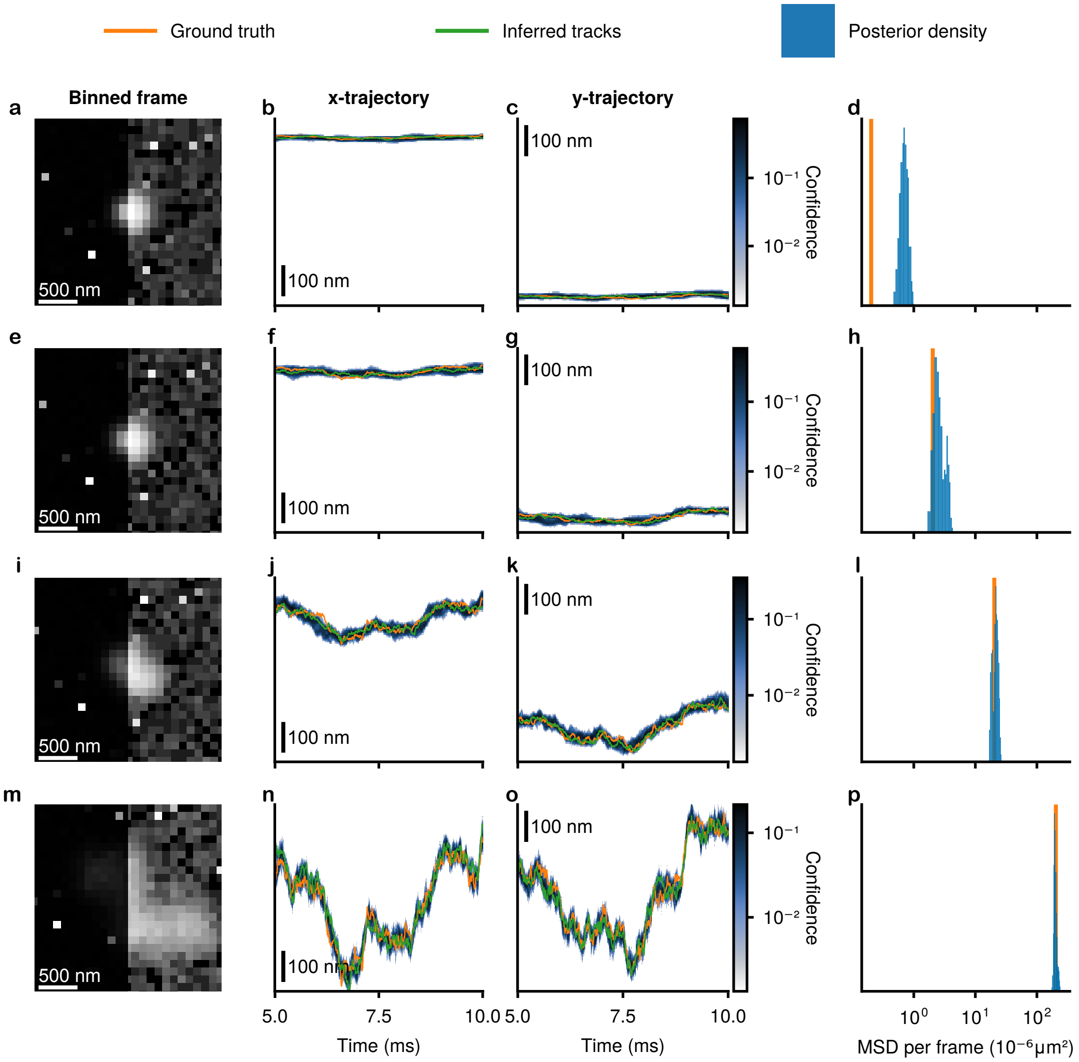


Figure S.3: Performance of SP$^2$T under variable diffusion coefficients (0.01 $\mu$m$^2$/s to 10 $\mu$m$^2$/s) using simulated data. Simulations assume a numerical aperture of 1.45, a refractive index of 1.515, an emission wavelength of 665 nm, and 10 $\mu$s exposure across 2,550 1-bit frames. The background noise is taken directly from Fig. 2 in the main text. **a**, Summed image of all 2,550 1-bit frames with a diffusion coefficient of 0.01 $\mu$m$^2$/s, for illustrative purposes. The left half is displayed with a linear colormap; the right half uses a logarithmic colormap to enhance low-intensity regions. **b–c**, Tracking results in both $x$- and $y$-directions. Ground truth tracks are shown in orange, while SP$^2$T's MAP tracks are in green. Shading reflects the confidence (posterior probability) of the emitter's position within each 10 nm bin; darker shades indicate higher confidence. **d**, Posterior distribution of SP$^2$T's per-frame MSD estimates, with the ground-truth value shown in orange. For such slow diffusion, SP$^2$T struggles to recover the correct MSD. **e–h**, Same layout as **a–d**, but with a diffusion coefficient of 0.1 $\mu$m$^2$/s. **i–l**, Same layout, with a diffusion coefficient of 1 $\mu$m$^2$/s. **m–p**, Same layout, with a diffusion coefficient of 10 $\mu$m$^2$/s.

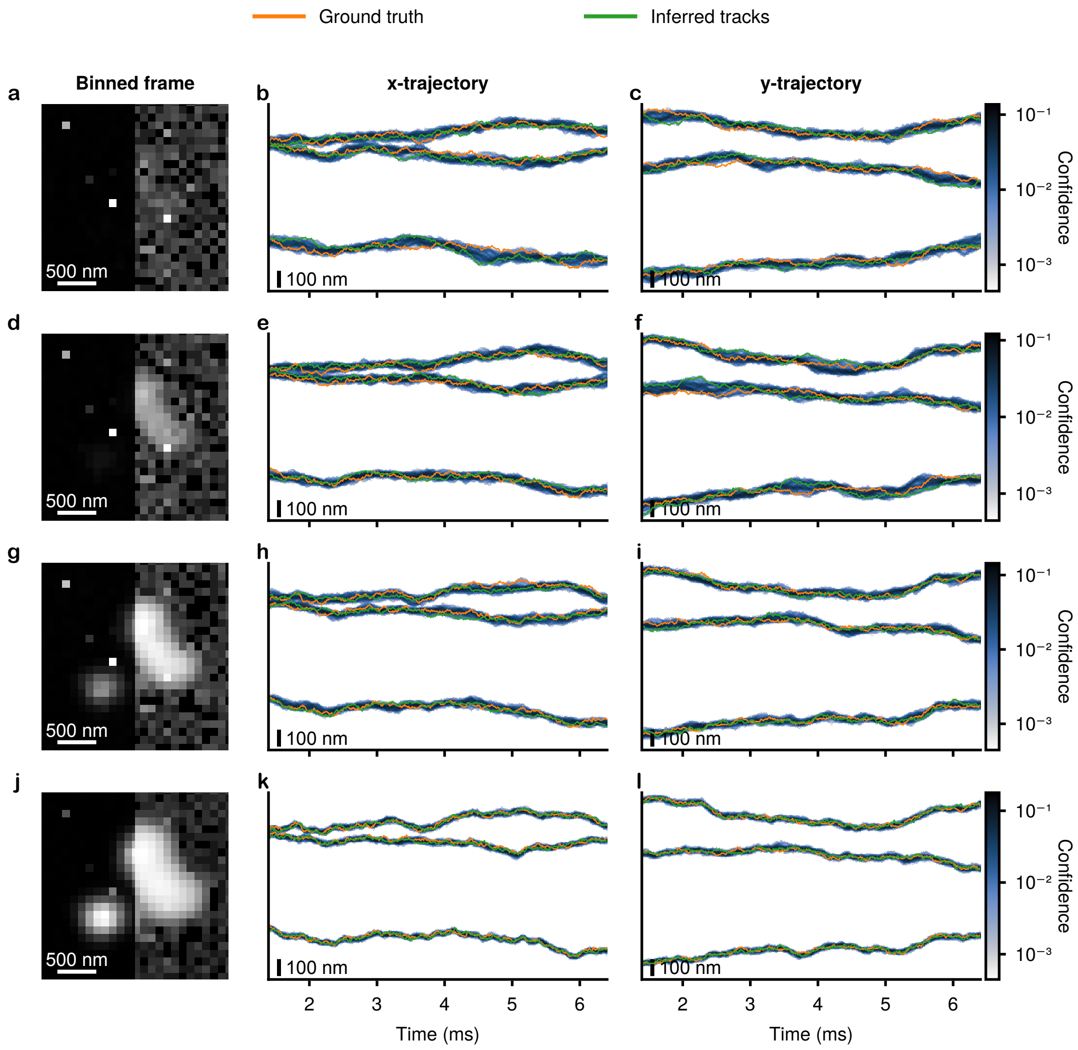


Figure S.4: Performance of SP$^2$T under varying emitter brightness levels ($4 \times 10^2\,\mathrm{s}^{-1}$ to $4 \times 10^5\,\mathrm{s}^{-1}$) using simulated data. A brightness of $4 \times 10^4\,\mathrm{s}^{-1}$ approximates the Cy3B dye used in Fig. 4 of the main manuscript. Simulations assume a numerical aperture of 1.45, refractive index 1.515, emission wavelength 665 nm, and 10 $\mu$s exposure across 2,550 1-bit frames. Three particles diffuse at 1 $\mu\mathrm{m}^2$/s. **a**, Summed image of all 2,550 1-bit frames with emitter brightness of $4\times10^2\,\mathrm{s}^{-1}$, for illustrative purposes. The left half uses a linear colormap; the right half uses a logarithmic colormap to enhance low-intensity detail. **b–c**, Tracking results in the $x$- and $y$-directions, respectively. Ground truth tracks are shown in orange, and in green, SP$^2$T's maximum a posteriori (MAP) tracks. Shading indicates posterior confidence (the probability of the emitter being located within a 10 nm bin); darker colors reflect higher confidence. **d–f**, Same layout as **a–c**, but for brightness $4 \times 10^3\,\mathrm{s}^{-1}$. **g–i**, Same layout for brightness $4 \times 10^4\,\mathrm{s}^{-1}$. **j–l**, Same layout for brightness $4 \times 10^5\,\mathrm{s}^{-1}$.

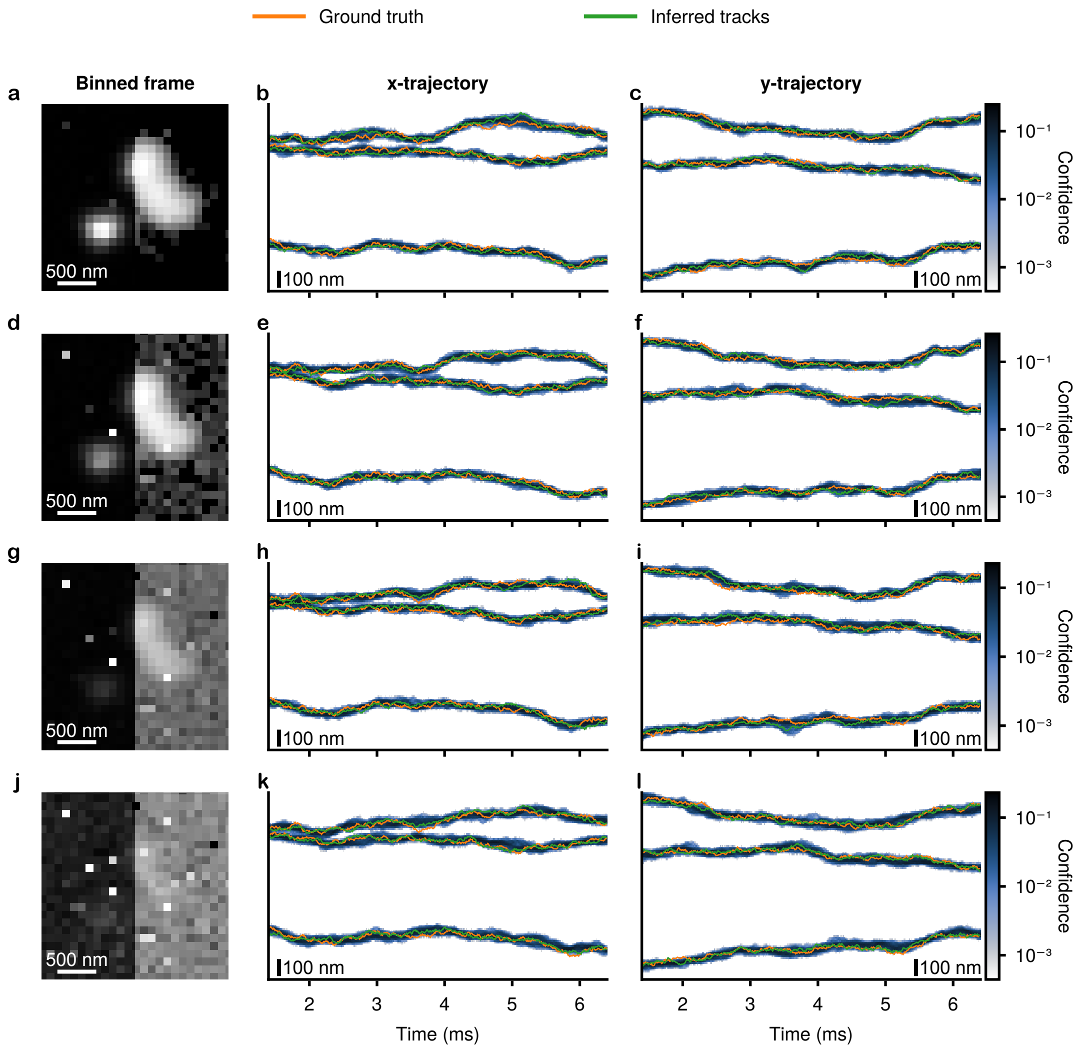


Figure S.5: Performance of SP$^2$T under varying dark count levels (0.1 to 100 signals per-frame per-pixel) using simulated data. A dark count level of 1 reflects realistic values as observed in Fig. 2a and Fig. 4 of the main text. Other values represent multiplicative factors relative to this baseline. Simulations assume a numerical aperture of 1.45, a refractive index of 1.515, an emission wavelength of 665 nm, and 10 $\mu$s exposure across 2,550 1-bit frames. Three particles are simulated to diffuse at 1 $\mu$m$^2$/s. **a**, Summed image of all 2,550 1-bit frames with a dark count of 0.1. The left half uses a linear colormap; the right half uses a logarithmic colormap to enhance low-intensity detail. **b–c**, Tracking results in the $x$- and $y$-directions, respectively. Ground truth tracks are shown in orange, and in green, SP$^2$T's maximum a posteriori (MAP) tracks. Shading indicates posterior confidence (the probability of the emitter being located within a 10 nm bin); darker colors reflect higher confidence. **d–f**, Same layout as **a–c**, but for dark count 1 s$^{-1}$. **g–i**, Same layout for dark count 10 s$^{-1}$. **j–l**, Same layout for dark count 100 s$^{-1}$.

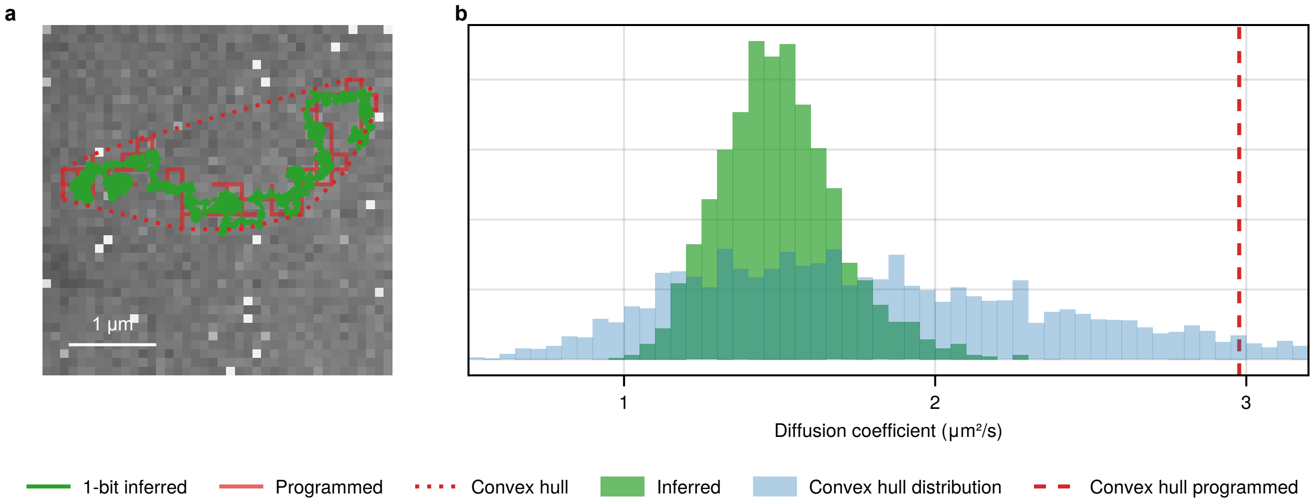


Figure S.6: Diffusion coefficient inference for surface-immobilized fluorescent dyes on a moving piezo stage shown in Fig. 2. Due to the fast, straight-line piezo stage movements, the diffusion coefficient is not particularly well defined and thus challenging to use to verify $SP^2T$'s accuracy. Here, we demonstrate a high-level validation that shows the inferred values to be reasonable. **a**, The 120 step programmed trajectory in red is locally not diffusive, but if considered from a coarse "area covered over time" perspective can yield an approximate diffusion coefficient by calculating the convex hull area $A_{\text{hull}}$ (red dotted boundary), and dividing by $\pi\Delta T$, where $\Delta T$ is the total time elapsed(*2*) **b**, Because diffusion coefficient is best understood as an ensemble quantity (expectation over trajectories), we can estimate the expected diffusion coefficient distribution (blue histogram) for the x/y step random walk by calculating the convex hull diffusion coefficient estimate for many realizations of a 120 step programmed trajectory, with the value estimated from the actual programmed trajectory considered marked by the red dashed line. The $SP^2T$ inferred posterior is shown in green. This is intended as a rough estimate to show that the obtained diffusion coefficient values are not unreasonable, even given the non-diffusive piezo stage motion. Experimental parameters include: exposure time of 40 $\mu$s, pixel size of 100 nm, numerical aperture of 1.49, refractive index of 1.52, and emission wavelength of 665 nm.

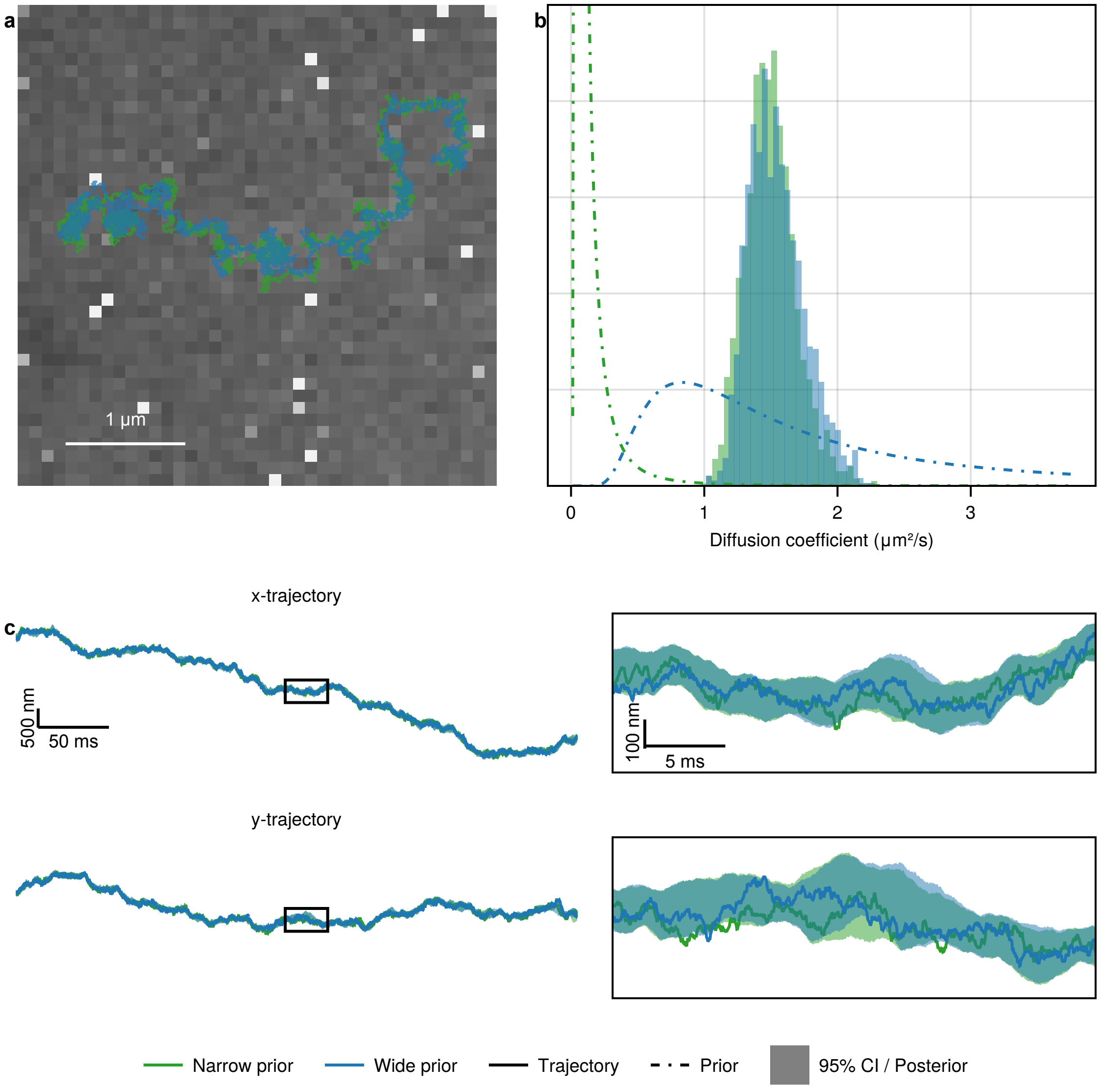


Figure S.7: **a**, Inference of particle tracks for programmed stage movement of fixed fluorescent dyes shown in Fig. 2 with two different diffusion coefficient prior choices. The MAP inferred trajectory for each case is plotted on the contrast-adjusted sum of all frames in the dataset. **b**, The form of the diffusion coefficient prior is inverse gamma, with support on $(0, \infty)$, with the more constraining prior shown in green with shape and scale parameters $[2,\ 1 \times 10^{-5}]$ and the wider prior in blue with parameters $[2,\ 2 \times 10^{-4}]$. The diffusion coefficient posteriors for both cases mostly overlap, with the blue having a slightly larger mean. **c**, Inferred x and y trajectories with 95% confidence intervals. In the zoomed-in regions, we can see that the confidence intervals largely overlap with minor variation. Overall, we see very little difference between the results for the two priors chosen. This is expected because the most important feature of a prior is its support, while the exact prior distribution can in general be overcome by the data, as is shown here, even for the narrow green prior offset from the posterior mass.

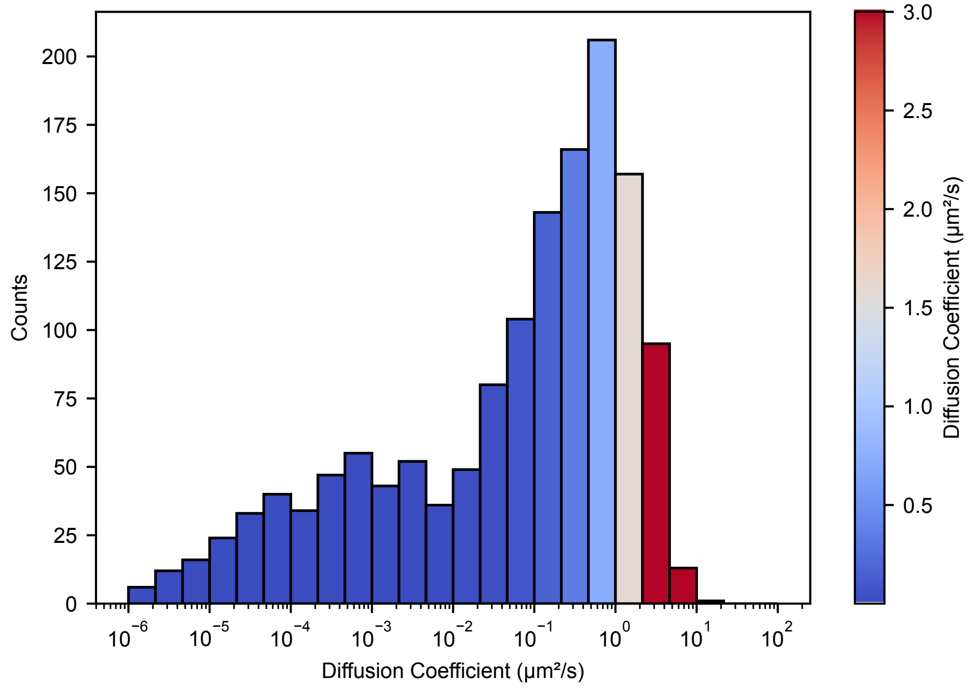


Figure S.8: Diffusion coefficient distribution obtained from EMCCD imaging. Analysis performed with TrackPy on 600 frames (512×512 pixels, 50 ms exposure) of the same sample shown in Fig. 3 (diffusing Atto565-labeled ganglioside GM1 molecules, highlighted in blue). Diffusion coefficients were estimated from mean-squared-displacement (MSD) fits.

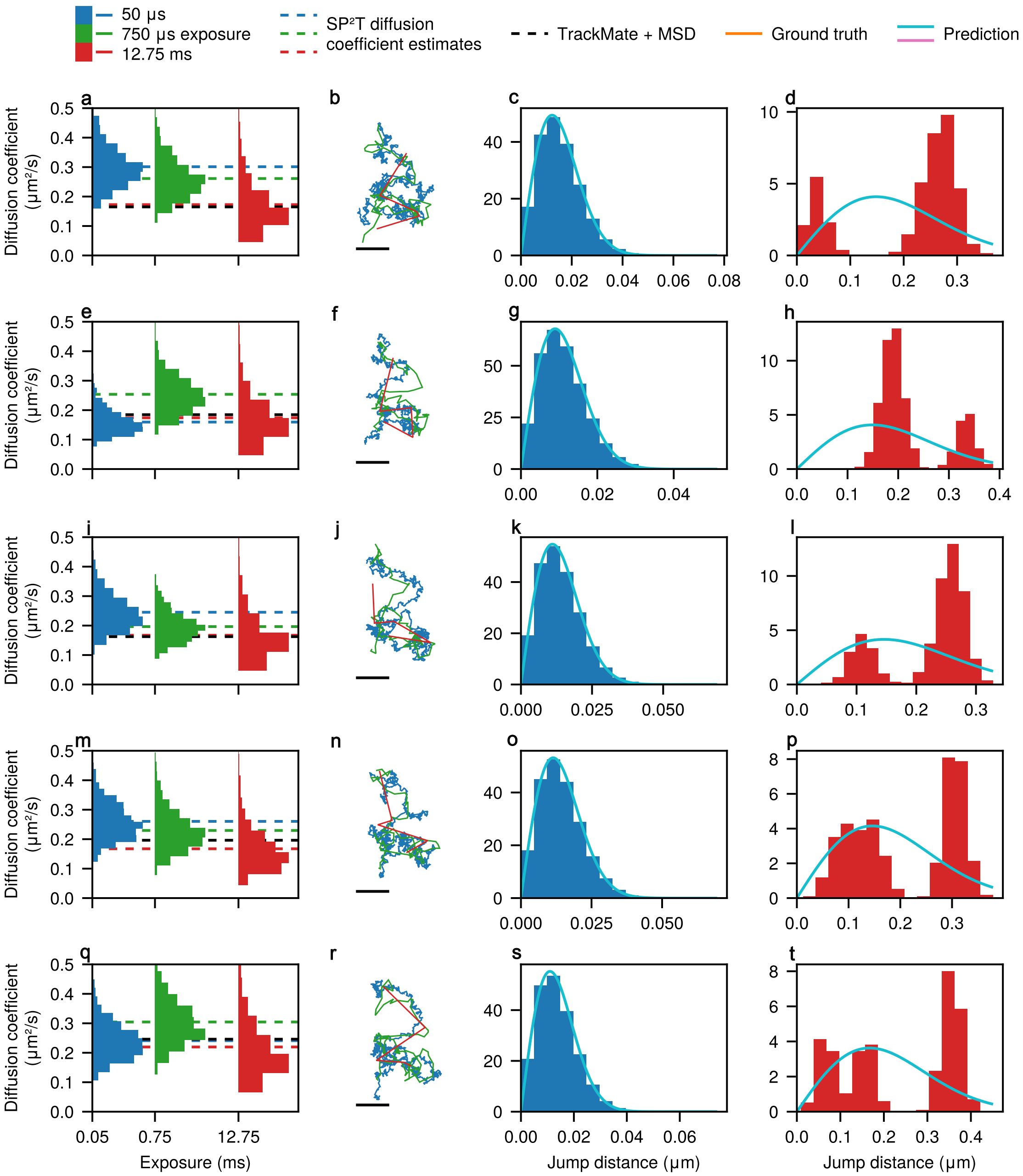


Figure S.9: Effect of camera exposure under pseudo-stroboscopic illumination. Diffusion coefficient estimates, particle trajectories, and jump-distance distributions are shown in the same layout as Fig. 3 of the main text, but with different frames retained in the stroboscopic scheme. **a–d** are identical to Figs. 3c to 3f, obtained when the stroboscopic scheme retains the first frame of every five-frame block. **e–h** show results when the second frame is retained; **i–l** when the third frame is retained; **m–p** when the fourth frame is retained; and **q–t** when the fifth frame is retained.

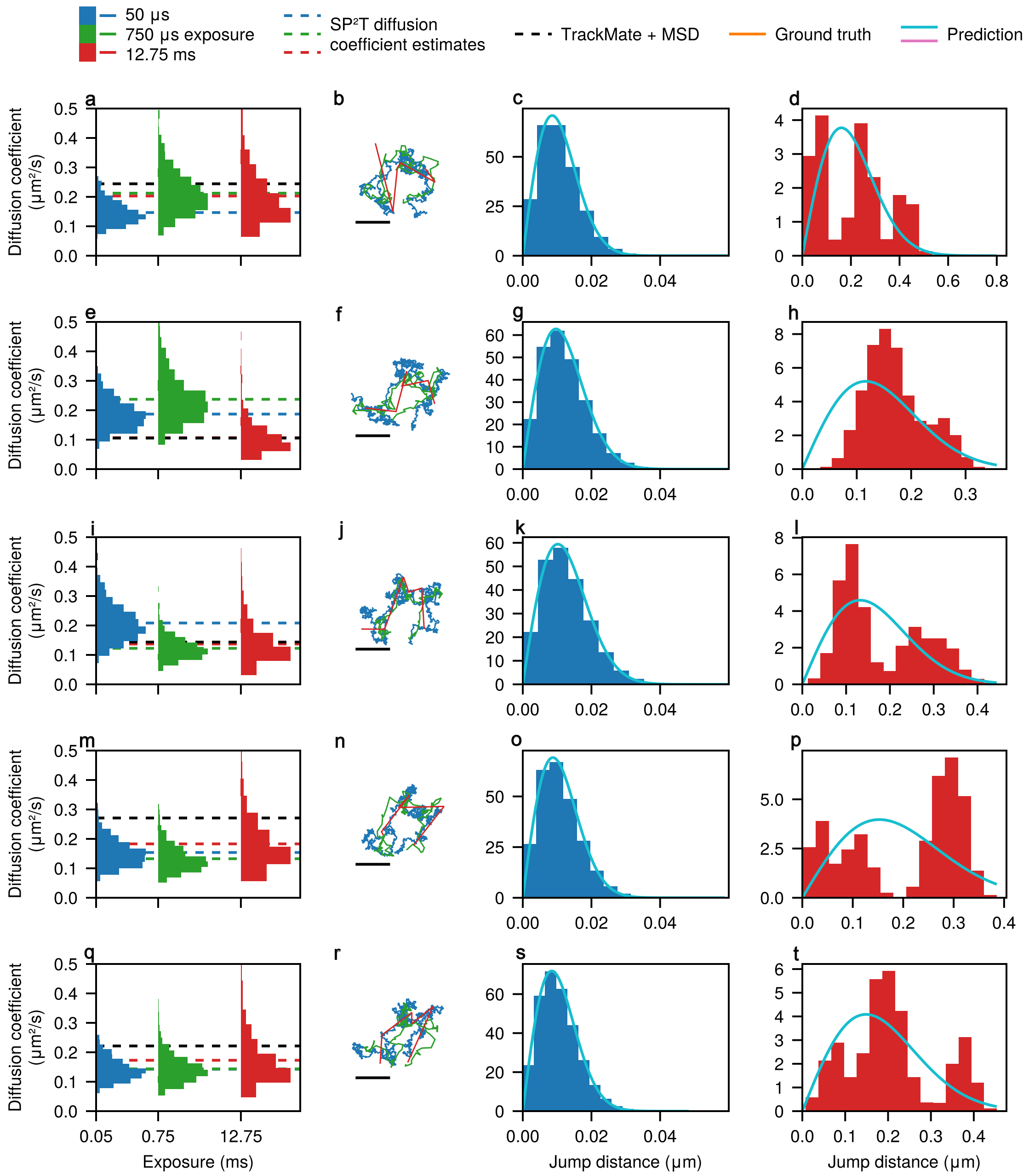


Figure S.10: Effect of camera exposure under pseudo-stroboscopic illumination. Diffusion coefficient estimates, particle trajectories, and jump-distance distributions are shown in the same layout as Fig. 3 of the main text, but taken from a different region of interest within the same dataset. **a–d** are identical to Figs. 3c to 3f, obtained when the stroboscopic scheme retains the first frame of every five-frame block. **e–h** show results when the second frame is retained; **i–l** when the third frame is retained; **m–p** when the fourth frame is retained; and **q–t** when the fifth frame is retained.

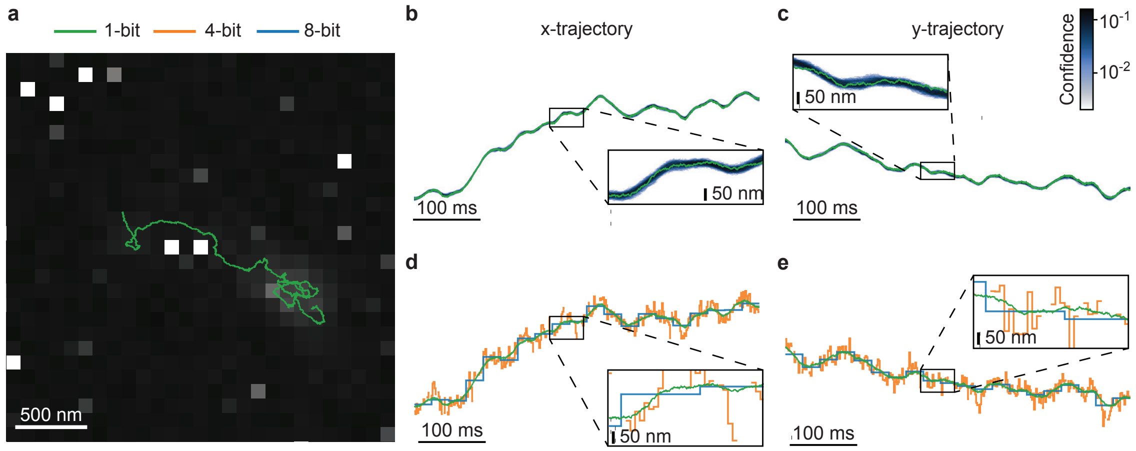


Figure S.11: Results for Cy3B-labeled aerolysin diffusing on a SLB for a 5,100 40×40 pixel 1-bit frames dataset, with the same experimental setup as Fig. 4. **a**, Example frame obtained by summing all 5,100 individual 40×40 1-bit frames zoomed in on this emitter, acquired at 100 $\mu$s exposure. The image has been contrast-adjusted for clarity, with the aerolysin track inferred by our framework overlaid. **b**, Confidence plot for the aerolysin's $x$-coordinate trajectory, where darker shading denotes higher confidence. The boxed region is magnified for closer inspection. **c**, Same as **b**, but for the $y$-coordinate trajectory. **d**, As ground truth is unavailable in real experiments, we approximate a control by comparing the aerolysin's 1-bit trajectory with 4-bit and 8-bit trajectories obtained using TrackMate(*1*). For clarity, we zoom in on the same region shown in **b**. Note the discontinuity in the 4-bit trajectory, which arises from TrackMate's failure to detect the particle under the low signal-to-noise ratio of the 4-bit frames. **e**, Same as **d**, but for the $y$-coordinate trajectory. Other experimental parameters include: numerical aperture of 1.49, refractive index of 1.518, and emission wavelength of 571 nm.

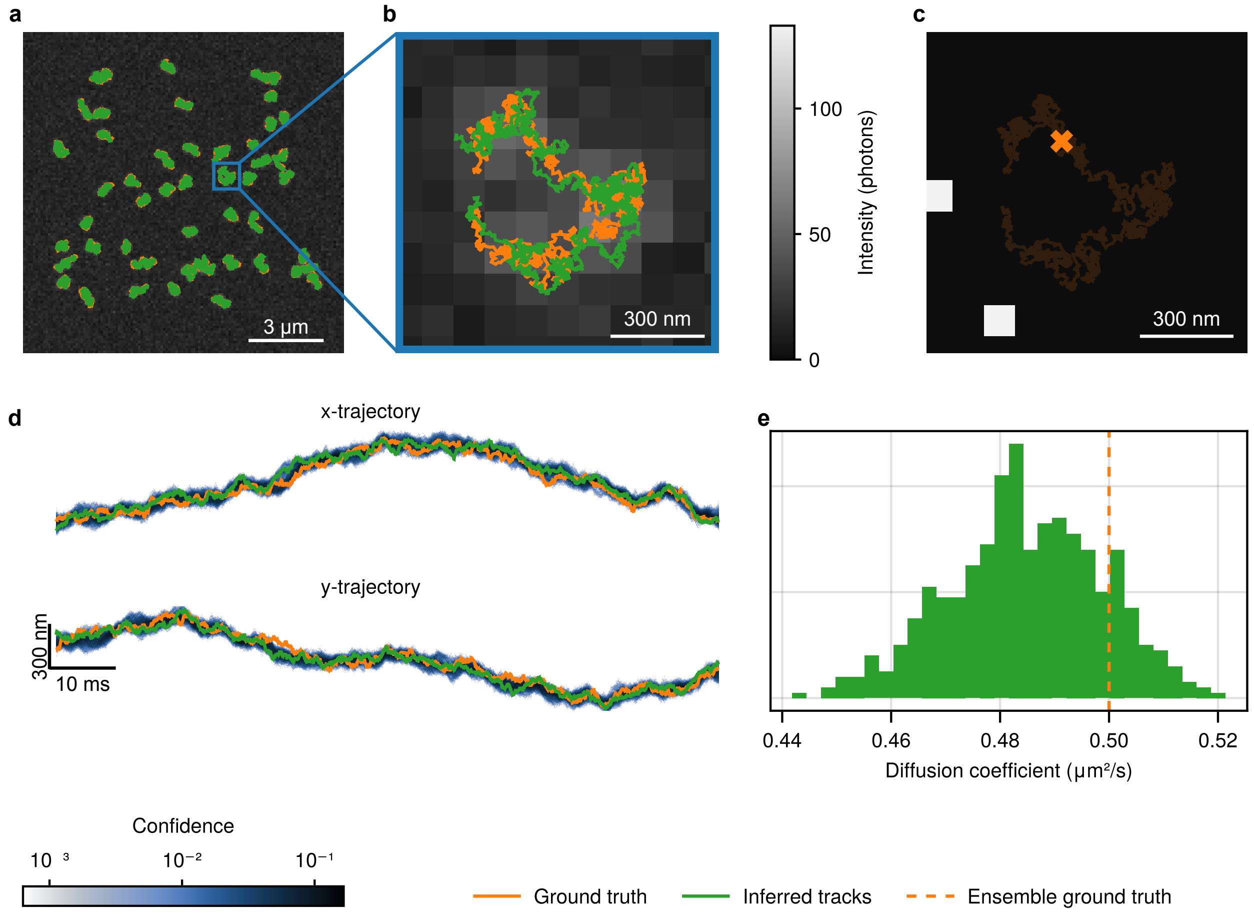


Figure S.12: Multiplexed tracking of 50 particles from 2000 50 $\mu$s 1-bit frames, similar to Fig. 5 tracking 100 particles. Initial tracks guess obtained by binning the ground truth trajectory with batch size 100, comparable to starting off with an initial track from applying Trackmate to binned frames. Inference was run parametrically (known number of particles) but when particle number is not known you can start by doing a short non-parametric run to refine the number of particles from the initial guess. **a**, Binned 128×128 frames of the simulation of 50 particles, for illustrative purposes, using an emitter brightness of 10,000 photons per second per emitter and a background photon rate of 14,000 photons per $\mu$m$^2$ per second to match the aerolysin data in Fig. 4, with ground truth and an inferred trajectory overlaid. **b**, Zoom-in to a single particle. **c**, Single photon frame showing two photon arrivals for the point in the trajectory indicated. **d**, Overlap of the ground truth trajectory and confidence region indicate excellent match between inferred and ground truth, with darker indicating greater confidence. **e**, The posterior distribution of inferred diffusion coefficient includes the true ensemble diffusion coefficient of 0.5 $\mu$m$^2$/s.

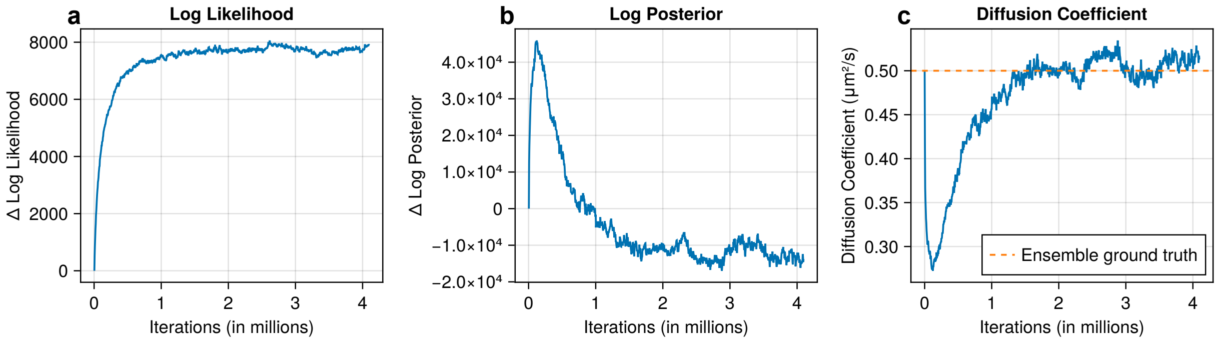


Figure S.13: Inference of 100 emitter tracks from 1000 100 $\mu$s 1-bit frames shown in Fig. 5 took 16 hours on an A30 GPU. **a**, $\Delta$log likelihood vs. iteration, **b**, $\Delta$ log posterior vs. iteration, and **c**, diffusion coefficient vs. iteration.

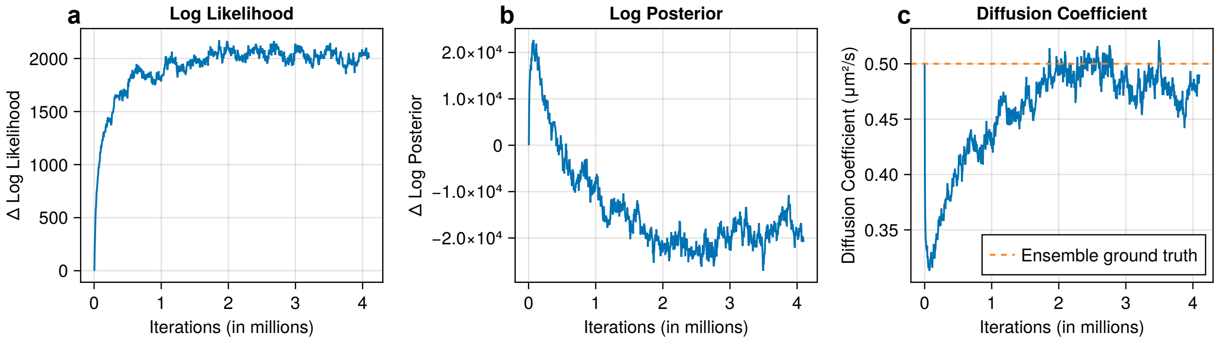


Figure S.14: Inference of 50 emitter tracks from 2000 50 $\mu$s 1-bit frames shown in Fig. S.12 took 22 hours on an A30 GPU. **a**, $\Delta$log likelihood vs. iteration, **b**, $\Delta$ log posterior vs. iteration, and **c**, diffusion coefficient vs. iteration.

### B Forward model

#### B.1 Observation model

We denote the total number of frames by $N$ and the total number of pixels by $P$. We use the indices $n$ and $p$ to represent the $n$th frame and the $p$th pixel, respectively. At the $n$th frame and the $p$th pixel, the raw measurement is denoted as $w_n^p$. Since a single-photon detector can only distinguish between the absence of photons (false, 0) and the presence of at least one photon (true, 1), the measurement $w_n^p$ must be sampled from a Bernoulli distribution. The parameter of this Bernoulli distribution is determined by the probability of detecting at least one photon, which is given by $1 - \text{Poisson}(0; u_n^p)$. Here, $u_n^p$ represents the incident intensity and $\text{Poisson}(0; u_n^p) = \exp(-u_n^p)$ represents the probability of detecting zero photons. Consequently, the probability of detecting at least one photon is $1 - \exp(-u_n^p)$, and the corresponding mathematical expression is:

$$w_n^p | u_n^p \sim \textbf{Bernoulli}\left(1 - \exp\left(-u_n^p\right)\right). \tag{S.1}$$

In order to model the incident photon counts $u_n^p$, we need to integrate the incident photon flux over the entire exposure time and pixel area. The exposure time interval in the $n$th frame is denoted as $[t_{n-1}, t_n]$, and the $p$th pixel is defined as the rectangular region $[x_{\text{min}}^p, x_{\text{max}}^p] \times [y_{\text{min}}^p, y_{\text{max}}^p]$. Therefore, the incident photon count in the $n$th frame and $p$th pixel can be represented by

$$u_n^p = \int_{x_{\text{min}}^p}^{x_{\text{max}}^p} \mathrm{d}x^p \int_{y_{\text{min}}^p}^{y_{\text{max}}^p} \mathrm{d}y^p \int_{t_{n-1}}^{t_n} \mathrm{d}t \, U(x^p, y^p, t), \tag{S.2}$$

where $U(x^p, y^p, t)$ is the incident photon flux with units of number per unit time per unit area. Additionally, the incident photon flux can be separated into the background and actual emitter (fluorescently labeled molecules) contributions. Thus, we can represent the incident photon flux as shown,

$$U\left(x^p, y^p, t\right) = U_{\text{back}}\left(x^p, y^p, t\right) + U_{\text{fluor}}\left(x^p, y^p, t\right), \tag{S.3}$$

where $U_{\text{back}}\left(x^p, y^p, t\right)$ is the background photon flux and $U_{\text{fluor}}\left(x^p, y^p, t\right)$ is the fluorescence photon flux. We assume that the background photon flux is constant over both space and time, so the function $U_{\text{back}}\left(x^p, y^p, t\right)$ can be simplified to just $F$. As for the fluorescence photon flux $U_{\text{fluor}}$, we model it as the sum of the individual emitter contributions

$$U_{\text{fluor}}\left(x^p, y^p, t\right) = \sum_{m=1}^{M} b^m U_{\text{fluor}}^m\left(x^p, y^p; \mathbf{x}^m(t)\right), \tag{S.4}$$

where $M$ stands for the total number of active loads, $b^m \in \{0, 1\}$ is a label (called load) for whether the $m$th emitter is active, and $\mathbf{x}^m(t)$ is the spatial location of the $m$th emitter at time $t$. The individual emitter contribution is given by

$$U_{\text{fluor}}^m\left(x^p, y^p; \mathbf{x}^m(t)\right) = h \, \text{PSF}\left(x^p, y^p; \mathbf{x}^m(t)\right). \tag{S.5}$$

In this equation, $h$ is the emission rate or brightness of the emitter, assumed constant across all emitters. The point spread function (PSF), denoted as $\mathrm{PSF}\left(x^p, y^p; \mathbf{x}\right)$, satisfies the condition $\int_{-\infty}^{\infty}\int_{-\infty}^{\infty} \mathrm{d}x^p \mathrm{d}y^p \mathrm{PSF}\left(x^p, y^p; \mathbf{x}^m(t)\right) = 1$. When these elements are plugged into the incident photon count formulation (equation (S.2)), we obtain

$$u_n^p = F^p A\tau + h \sum_{m=1}^{M} b^m \int_{t_{n-1}}^{t_n} \mathrm{d}t \int_{x_{\min}^p}^{x_{\max}^p} \mathrm{d}x^p \int_{y_{\min}^p}^{y_{\max}^p} \mathrm{d}y^p \mathrm{PSF}\left(x^p, y^p; \mathbf{x}^m(t)\right). \tag{S.6}$$

Here, $A^p = (x_{\max}^p - x_{\min}^p) \times (y_{\max}^p - y_{\min}^p)$ represents the area of the $p$th pixel, $t_n$ is the time when the $n$th frame is produced, and $\tau$ is the exposure time of each frame. We note that $\tau \le t_n - t_{n-1}$ due to camera dead time, and we assume no gaps between pixels, so $x_{\max}^p = x_{\min}^{p+1}$, $y_{\max}^p = y_{\min}^{p+1}$.

To evaluate the triple integral of equation (S.6), we must specify the integrand, namely the PSF. In practice, we can use any pre-calibrated form, including forms induced by optical aberration. For concreteness here, we use the circular Gaussian Lorentzian PSF given by

$$\mathrm{PSF}\left(x^p, y^p; \mathbf{x}^m(t)\right) = \frac{1}{2\pi\sigma_{z^m}^2} \exp\left(-\frac{\left(x^p - x^m\right)^2 + \left(y^p - y^m\right)^2}{2\sigma_{z^m}^2}\right), \tag{S.7}$$

$$\sigma_{z^m}^2 = \sigma_{\mathrm{ref}}^2 \left[1 + \left(\frac{z^m}{Z_{\mathrm{ref}}}\right)^2\right], \quad Z_{\mathrm{ref}} = \frac{4\pi n_{\mathrm{fluid}}}{\lambda} \sigma_{\mathrm{ref}}^2. \tag{S.8}$$

The PSF's width, denoted by $\sigma_{z^m}$, is given in equation (S.8), where $n_{\mathrm{fluid}}$ represents the refractive index of the immersion media in which the objective lens and the specimen are immersed, and $\lambda$ is the vacuum wavelength of the emitted photons. The lateral Gaussian parameter $\sigma_{\mathrm{ref}}$ is calculated differently based on the fluorescence microscope model(*3*). In this work, we consider the nonparaxial widefield fluorescence microscope(*4*) and $\sigma_{\mathrm{ref}}$ is given by

$$\sigma_{\mathrm{ref}} = \frac{\lambda}{2\pi n_{\mathrm{fluid}}} \sqrt{\frac{7\left(1 - \cos^{3/2}\alpha\right)}{4 - 7\cos^{3/2}\alpha + 3\cos^{7/2}\alpha}} \tag{S.9}$$

where $\alpha$ is the maximal convergence semi-angle of the objective.

Now that the PSF has been specified, we can evaluate the integral in equation (S.6) analytically. The area integral is then,

$$\begin{aligned} &\int_{x_{\min}^p}^{x_{\max}^p} \mathrm{d}x^p \int_{y_{\min}^p}^{y_{\max}^p} \mathrm{d}y^p \, \mathrm{PSF}\left(x^p, y^p; \mathbf{x}^m(t)\right) \\ &= \frac{1}{4}\left[\mathbf{erf}\left(\frac{x_{\max}^p - x^m}{\sigma_{z^m}\sqrt{2}}\right) - \mathbf{erf}\left(\frac{x_{\min}^p - x^m}{\sigma_{z^m}\sqrt{2}}\right)\right]\left[\mathbf{erf}\left(\frac{y_{\max}^p - y^m}{\sigma_{z^m}\sqrt{2}}\right) - \mathbf{erf}\left(\frac{y_{\min}^p - y^m}{\sigma_{z^m}\sqrt{2}}\right)\right]. \end{aligned} \tag{S.10}$$

It is worth noting that as we consider single-photon detectors with very high (97.7 kHz) data acquisition rates, the motion blur of emitters becomes negligible for most particles in biological systems(*5*). Therefore, with $g_n^{m,p}$ representing the result of the integral above, we have

$$u_n^p = F^p + h \sum_{m=1}^{M} b^m g_n^{m,p}. \tag{S.11}$$

#### B.2 Markovian dynamics for spatial trajectories

In equation (S.11), the pixel area $A^p$, exposure time $\tau$, dark counts $F$, emission rate $h$, and load $b^m$ are input parameters in the data synthesis process. The remaining task is to sample each emitter's spatial trajectory and compute $g_n^{m,p}$. In this study, we only consider a single diffusive species. Therefore, the motion model with a constant mean squared displacement MSD, can be represented by

$$\mathbf{x}_n^m | \mathbf{x}_{n-1}^m, \text{MSD} \sim \mathbf{Normal}_3 \left( \mathbf{x}_{n-1}^m, \text{MSD}\, \mathbf{I} \right). \tag{S.12}$$

Here, $\mathbf{I}$ is the three-dimensional identity matrix. The initial positions of each emitter are sampled from Normal distributions with fixed means and variances as follows

$$x_1^m \sim \mathbf{Normal} \left( \mu_{xy}, \sigma_{xy}^2 \right), \tag{S.13}$$

$$y_1^m \sim \mathbf{Normal} \left( \mu_{xy}, \sigma_{xy}^2 \right), \tag{S.14}$$

$$z_1^m \sim \mathbf{Normal} \left( \mu_{z}, \sigma_{z}^2 \right). \tag{S.15}$$

#### B.3 Model summary

The entire procedure outlined in section B can be summarized in Fig. S.15. For ease of reference, we have also included a list of all key equations and probability distributions discussed up to this point in the text

$$x_1^m \sim \mathbf{Normal} \left( \mu_{xy}, \sigma_{xy}^2 \right), \tag{S.16}$$

$$y_1^m \sim \mathbf{Normal} \left( \mu_{xy}, \sigma_{xy}^2 \right), \tag{S.17}$$

$$z_1^m \sim \mathbf{Normal} \left( \mu_{z}, \sigma_{z}^2 \right), \tag{S.18}$$

$$\mathbf{x}_n^m \middle| \mathbf{x}_{n-1}^m, \text{MSD} \sim \mathbf{Normal}_3 \left( \mathbf{x}_{n-1}^m, \text{MSD}\, \mathbf{I} \right) \tag{S.19}$$

$$g_n^{m,p} = \frac{1}{4} \left[ \mathbf{erf} \left( \frac{x_{\text{max}}^p - x_n^m}{\sigma_{z_n^m} \sqrt{2}} \right) - \mathbf{erf} \left( \frac{x_{\text{min}}^p - x_n^m}{\sigma_{z_n^m} \sqrt{2}} \right) \right] \times \left[ \mathbf{erf} \left( \frac{y_{\text{max}}^p - y_n^m}{\sigma_{z_n^m} \sqrt{2}} \right) - \mathbf{erf} \left( \frac{y_{\text{min}}^p - y_n^m}{\sigma_{z_n^m} \sqrt{2}} \right) \right], \tag{S.20}$$

$$u_n^p = F^p + h\tau \sum_{m=1}^{M} g_n^{m,p}, \tag{S.21}$$

$$w_n^p | u_n^p \sim \mathbf{Bernoulli} \left( 1 - \exp\left( -u_n^p \right) \right). \tag{S.22}$$

Videos can now be synthesized by following these steps:

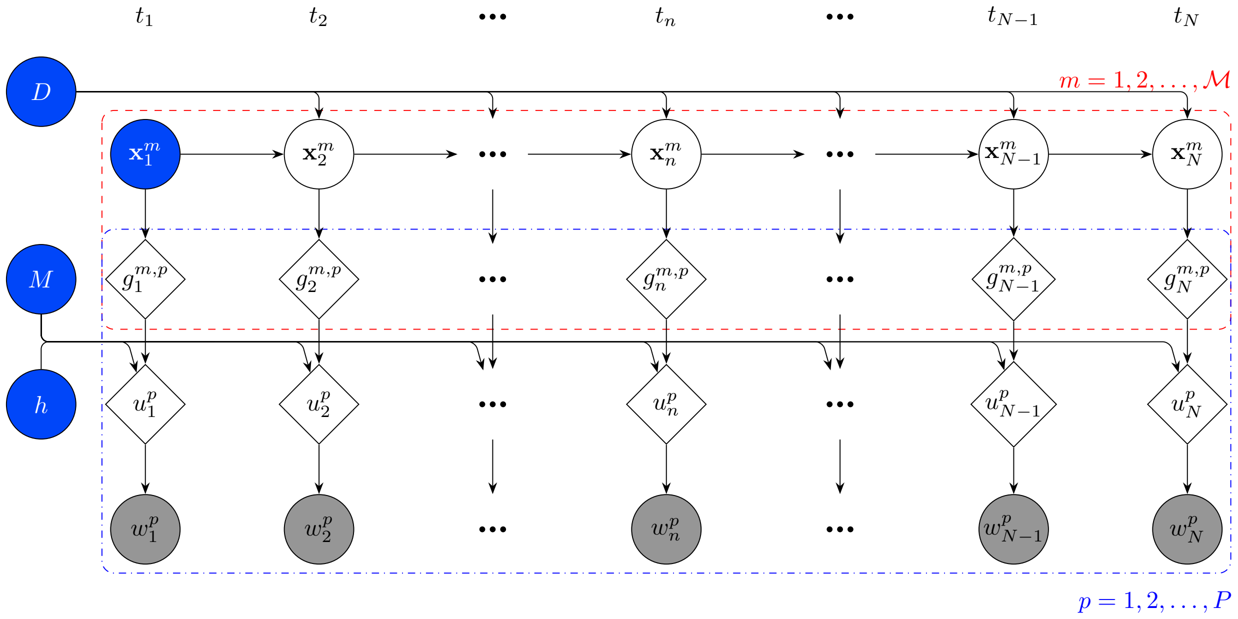


Figure S.15: Graphic model of our tracking framework.

1. Specify all camera parameters;
2. Sample the initial position of each emitter using equations (S.16) and (S.18);
3. Determine the subsequent spatial trajectory of each emitter using equation (S.19);
4. Calculate the photon number $u_n^p$ for each pixel and each frame using equation (S.21);
5. Sample from equation (S.22) to generate actual measurements.

A note on this data generation is in order. Motion blur artifacts will naturally appear because of step 4 as we evaluate equation (S.20). Other features, such as emitters moving out of focus and their spots becoming broader and blurrier, are also incorporated as follows from equation (S.20).

### C Inference

The main objective of SP$^2$T is to learn spatial trajectories of all emitters, $\mathbf{x}_{1:N}^{1:M}$, as well as their loads $b^{1:M}$, and the MSD, where $\mathbf{x}_{1:N}^{1:M} \equiv \{\mathbf{x}_n^m | n = 1, \ldots, N, m = 1, \ldots, M\}$, and $b^{1:M} \equiv \{b^m | m = 1, \ldots, M\}$. In the Bayesian paradigm, estimating these quantities is equivalent to sampling from the posterior probability distribution $\mathcal{P}\left(b^{1:M}, \mathbf{x}_{1:N}^{1:M}, \text{MSD} \middle| w_{1:N}^{1:P}\right)$ where $w_{1:N}^{1:P} \equiv \{w_n^p | n = 1, \ldots, N, p = 1, \ldots, P\}$ represents all the measurements. The rest of this section will provide a detailed description of how to sample from this posterior distribution.

#### C.1 Posterior construction

The complete posterior probability distribution is represented in a specific format, derived using Bayes' theorem:

$$\mathcal{P}\left(b^{1:M}, \mathbf{x}_{1:N}^{1:M}, \text{MSD} \middle| w_{1:N}^{1:P}\right) = \frac{\mathcal{P}\left(w_{1:N}^{1:P} \middle| b^{1:M}, \mathbf{x}_{1:N}^{1:M}, \text{MSD}\right) \mathcal{P}\left(b^{1:M}, \mathbf{x}_{1:N}^{1:M}, \text{MSD}\right)}{\mathcal{P}\left(w_{1:N}^{1:P}\right)}. \tag{S.23}$$

The term $\mathcal{P}\left(w_{1:N}^{1:P}\right)$ in the denominator is recognized as the evidence. Since it is independent of the quantities of interest, $b^{1:M}$, $\mathbf{x}_{1:N}^{1:M}$, and MSD, we can treat $\mathcal{P}\left(w_{1:N}^{1:P}\right)$ as merely a normalization constant.

The first term in the numerator of equation (S.23), $\mathcal{P}\left(w_{1:N}^{1:P} \middle| b^{1:M}, \mathbf{x}_{1:N}^{1:M}, D\right)$, termed the observation likelihood, can be determined using equations (S.20) to (S.22). It is important to note that, conditioned on $b^{1:M}$ and $\mathbf{x}_{1:N}^{1:M}$, the MSD does not influence the measurements. Therefore, we can simplify the observation likelihood expression to $\mathcal{P}\left(w_{1:N}^{1:P} \middle| b^{1:M}, \mathbf{x}_{1:N}^{1:M}\right)$. Given the frequency of this likelihood's usage in the text, we will abbreviate it as $\mathcal{L}$ for notational ease. We also assume that pixels of a camera provide independent measurements, and measurements from the same pixel are not temporally correlated. This leads to the following equations:

$$\begin{aligned}
\mathcal{L} &= \prod_{n=1}^{N} \prod_{p=1}^{P} \mathcal{P}\left(w_n^p \middle| b^{1:M}, \mathbf{x}_n^{1:M}\right) = \prod_{n=1}^{N} \prod_{p=1}^{P} \textbf{Bernoulli}\left(w_n^p; 1 - \exp\left(-u_n^p\right)\right) \\
&= \prod_{n=1}^{N} \prod_{p=1}^{P} \left[1 - \exp\left(-u_n^p\right)\right]^{w_n^p} \exp\left[-\left(1 - w_n^p\right) u_n^p\right] \\
&= \prod_{n=1}^{N} \prod_{p=1}^{P} \left[\exp\left(u_n^p\right) - 1\right]^{w_n^p} \exp\left(-u_n^p\right), \qquad \text{(S.24)} \\
\ln \mathcal{L} &= \sum_{n=1}^{N} \sum_{p=1}^{P} w_n^p \ln\left[\exp\left(u_n^p\right) - 1\right] - u_n^p. \qquad \text{(S.25)}
\end{aligned}$$

The second term in the numerator of equation (S.23), $\mathcal{P}\left(b^{1:M}, \mathbf{x}_{1:N}^{1:M}, \text{MSD}\right)$, is termed the prior within the Bayesian framework. A prior probability distribution encapsulates the pre-existing knowledge about its arguments. Given that the emitter's diffusion dynamics, brightness, load, and the system's dark counts are assumed to be mutually independent, we can express $\mathcal{P}\left(b^{1:M}, \mathbf{x}_{1:N}^{1:M}, \text{MSD}\right)$ as the product of several individual priors:

$$\mathcal{P}\left(b^{1:M}, \mathbf{x}_{1:N}^{1:M}, \text{MSD}\right) = \mathcal{P}\left(b^{1:M}\right) \mathcal{P}\left(\mathbf{x}_{1:N}^{1:M}, \text{MSD}\right). \tag{S.26}$$

The specific choice for each prior will be detailed in the following sections.

By combining equations (S.23) and (S.26), we obtain the following expression

$$\mathcal{P}\left(b^{1:M}, \mathbf{x}_{1:N}^{1:M}, \text{MSD} \middle| w_{1:N}^{1:P}\right) \propto \mathcal{P}\left(b^{1:M}\right) \mathcal{P}\left(\mathbf{x}_{1:N}^{1:M}, \text{MSD}\right) \mathcal{L} \tag{S.27}$$

which coincides with the posterior probability distribution from which we sample.

While this distribution is too complex to allow for direct sampling, we use a Gibbs sampling scheme to update the parameters iteratively from their conditional posteriors. Specifically, the sampling procedure follows this sequence

1. Initial values for $\mathbf{x}_{1:N}^{1:M}$, $b^{1:M}$, MSD, are set by hand.
2. $b^{1:M}$ is updated by sampling from
$$\mathcal{P}\left(b^{1:M}\middle|w_{1:N}^{1:P}, \mathbf{x}_{1:N}^{1:M}, D\right) \propto \mathcal{P}\left(b^{1:M}\right)\mathcal{L}; \tag{S.28}$$
3. $\mathbf{x}_{1:N}^{1:M}$ and MSD are updated by sampling from
$$\mathcal{P}\left(\mathbf{x}_{1:N}^{1:M}, D\middle|w_{1:N}^{1:P}, b^{1:M}\right); \tag{S.29}$$
4. $\mathbf{x}_{1:N}^{\mathcal{M}_1}$ is updated again by sampling from
$$\mathcal{P}\left(\mathbf{x}_{1:N}^{\mathcal{M}_1}\middle|w_{1:N}^{1:P}, b^{1:M}, \mathbf{x}_{1:N}^{\mathcal{M}_0}, D\right) \propto \mathcal{P}\left(\mathbf{x}_{1:N}^{\mathcal{M}_1}\middle|D\right)\mathcal{P}\left(w_{1:N}^{1:P}\middle|b^{1:M}, \mathbf{x}_{1:N}^{1:M}\right) \tag{S.30}$$
where $\mathcal{M}_i$ is the collection of all $m$'s with $b_m = i$;
5. Repeat steps 2 through 4.

#### C.2 Sampling loads

In the preceding sections, we have discussed how an emitter's load is a binary variable that labels whether the particle contributes to the observation likelihood. According to this definition, the loads of different emitters should be independent. Consequently, there are $2^M$ potential load configurations, and as $M$ increases, directly sampling emitter loads becomes computationally infeasible. Instead of direct sampling, only a small subset of all loads can be updated at each Gibbs iteration, which often leads to longer mixing time.

To allow for direct load sampling, even with a larger value of $M$, we can introduce a hierarchy between emitter loads to reduce the total number of possible configurations. Specifically, we define the complete load prior as follows:

$$\mathcal{P}\left(b^{1:M}\right) = \mathcal{P}\left(b^{1}\right)\prod_{m=2}^{M}\mathcal{P}\left(b^{m}\middle|b^{m-1}\right). \tag{S.31}$$

In this study, we select $\mathcal{P}(b^1)$ and $\mathcal{P}(b^m|b^{m-1})$ such that

$$\mathcal{P}\left(b^{1}\right) = q^{b^1}\left(1-q\right)^{1-b^1}, \tag{S.32}$$
$$\mathcal{P}\left(b^{m}\middle|b^{m-1}\right) = 1 + b^{m-1}\left[q^{b^m}\left(1-q\right)^{1-b^m} - 1\right]. \tag{S.33}$$

For clarity, the two equations above can be alternatively expressed as:

$$b^1 \sim \textbf{Bernoulli}\left(q\right), \tag{S.34}$$

$$b^m \middle| b^{m-1} \sim \begin{cases} \textbf{Bernoulli}\left(q\right), & b^{m-1} = 1, \\ \textbf{Bernoulli}\left(0\right), & b^{m-1} = 0. \end{cases} \tag{S.35}$$

By employing this approach, the load of emitter $m$ can only be activated when the load of emitter $m-1$ is also turned on. Consequently, this prior distribution for loads reduces the number of potential configurations from $2^M$ to $M+1$.

In addition to the prior probability distribution described above, we also independently shuffle the labels of both active and inactive emitters. This relabeling is necessary because particle labels serve only as dummy indices and carry no intrinsic meaning; allowing label switches ensures proper mixing and symmetry in the sampling process. This shuffling can be mathematically depicted as a bijective map, $\mathcal{S} : 1{:}M \to 1{:}M$, such that

$$\begin{cases} \mathcal{S}\left(m\right) \in 1{:}B, & m \in 1{:}B, \\ \mathcal{S}\left(m\right) \in \left(B+1\right){:}M, & m \in \left(B+1\right){:}M, \end{cases} \tag{S.36}$$

where $B = \sum_{m=1}^{M} b^m$.

It is crucial to emphasize that this shuffling does not change a sample's posterior. Hence, the shuffling can be viewed as transitioning between statistically equivalent samples within the sample space. Now, we can proceed to sample $b^{1:M}$ jointly from the target probability distribution specified in equation (S.28). Specifically, we further write equation (S.28) as

$$\mathcal{P}\left(b^{1:M} \middle| w_{1:N}^{1:P}, \mathbf{x}_{1:N}^{1:M}\right) \propto \mathcal{P}\left(b^{1:M}\right) \mathcal{P}\left(w_{1:N}^{1:P} \middle| b^{1:M}, \mathbf{x}_{1:N}^{1:M}\right). \tag{S.37}$$

The math remains the same for each load, so we will just demonstrate the general case of updating $b^m$. The individual posterior of $b^m$ is given by

$$\mathcal{P}\left(b^{m} \middle| w_{1:N}^{1:P}, \mathbf{x}_{1:N}^{1:M}\right) \propto \mathcal{P}\left(b^{m}\right) \mathcal{P}\left(w_{1:N}^{1:P} \middle| b^{1:M}, \mathbf{x}_{1:N}^{1:M}\right). \tag{S.38}$$

Then the probabilities of both possible values of $b^m$ are

$$\begin{aligned} \mathcal{P}_{\text{off}} &\equiv \mathcal{P}\left(b^m = 0 \middle| b^{1:m-1,m+1:M}, w_{1:N}^{1:P}, \mathbf{x}_{1:N}^{1:M}\right) \\ &= A\frac{M-\gamma}{M}\mathcal{L}\left(b^{1:m-1,m+1:M}, b^m = 0, \mathbf{x}_{1:N}^{1:M}\right), \end{aligned} \tag{S.39}$$

$$\begin{aligned} \mathcal{P}_{\text{on}} &\equiv \mathcal{P}\left(b^m = 1 \middle| b^{1:m-1,m+1:M}, w_{1:N}^{1:P}, \mathbf{x}_{1:N}^{1:M}\right) \\ &= A\frac{\gamma}{M}\mathcal{L}\left(b^{1:m-1,m+1:M}, b^m = 1, \mathbf{x}_{1:N}^{1:M}\right), \end{aligned} \tag{S.40}$$

where $A$ is a normalization constant. Therefore, each $b^m$ is updated according to

$$b^m \sim \textbf{Bernoulli}\left(\mathcal{P}_{\text{on}}\right). \tag{S.41}$$

It is important to consider numerical stability when implementing the load sampling scheme. As before, all operations are performed in logarithmic space. Most of the modifications are straightforward algebraic manipulations, but it is worth noting that equation (S.41) in logarithmic space can be sampled using the "Gumbel-max" trick(*6*). Specifically, let

$$g_{\text{on}} \sim \textbf{Gumbel}(0, 1) \tag{S.42}$$

and

$$g_{\text{off}} \sim \textbf{Gumbel}(0, 1) \tag{S.43}$$

then

$$b^m = \begin{cases} 0, & \text{if } g_{\text{on}} + \ln \mathcal{P}_{\text{on}} < g_{\text{off}} + \ln \mathcal{P}_{\text{off}}; \\ 1, & \text{if } g_{\text{on}} + \ln \mathcal{P}_{\text{on}} \geq g_{\text{off}} + \ln \mathcal{P}_{\text{off}}. \end{cases} \tag{S.44}$$

In this paragraph, we will demonstrate an example of updating two loads, $b^{m_1}$ and $b^{m_2}$ (where $m_2 > m_1$), simultaneously. The majority of the derivation in this section remains valid, and the posterior of $b^{m_1}$ and $b^{m_2}$ becomes a Categorical distribution. The probabilities for each category, $(0, 0)$, $(1, 0)$, $(0, 1)$, and $(1, 1)$, are given by

$$\mathcal{P}_{(0,0)} = A \frac{(M-\gamma)^2}{M^2} \mathcal{L}\left(b^{1:m_1-1, m_1+1:m_2-1, m_2+1:M}, b^{m_1} = 0, b^{m_2} = 0, \mathbf{x}_{1:N}^{1:M}\right), \tag{S.45}$$

$$\mathcal{P}_{(0,1)} = A \frac{\gamma(M-\gamma)}{M^2} \mathcal{L}\left(b^{1:m_1-1, m_1+1:m_2-1, m_2+1:M}, b^{m_1} = 0, b^{m_2} = 1, \mathbf{x}_{1:N}^{1:M}\right), \tag{S.46}$$

$$\mathcal{P}_{(1,0)} = A \frac{\gamma(M-\gamma)}{M^2} \mathcal{L}\left(b^{1:m_1-1, m_1+1:m_2-1, m_2+1:M}, b^{m_1} = 1, b^{m_2} = 0, \mathbf{x}_{1:N}^{1:M}\right), \tag{S.47}$$

$$\mathcal{P}_{(1,1)} = A \frac{\gamma^2}{M^2} \mathcal{L}\left(b^{1:m_1-1, m_1+1:m_2-1, m_2+1:M}, b^{m_1} = 1, b^{m_2} = 1, \mathbf{x}_{1:N}^{1:M}\right). \tag{S.48}$$

The "Gumbel-max" technique should also be modified accordingly in order to accommodate more categories. Specifically, we need to independently sample $g_{(0,0)}$, $g_{(0,1)}$, $g_{(1,0)}$, and $g_{(1,1)}$ and determine the category the maximizes $g + \ln \mathcal{P}$.

#### C.3 Sampling trajectories and diffusion coefficient

As previously mentioned in section C.1, we know that an emitter only contributes to observations when its load is active ($b^m = 1$). This also implies that the posterior probability distribution of the trajectory of emitters with active loads includes observations, while the distribution of emitters with inactive loads does not. Therefore, to sample from the target distribution $\mathcal{P}\left(\mathbf{x}_{1:N}^{1:M}, \text{MSD} \middle| w_{1:N}^{1:P}, b^{1:M}\right)$, we can update the trajectories of both active ($\mathbf{x}_{1:N}^{\mathcal{M}_1}$) and inactive ($\mathbf{x}_{1:N}^{\mathcal{M}_0}$) emitters separately. This results in the following posterior form

$$\mathcal{P}\left(\mathbf{x}_{1:N}^{1:M}, \text{MSD} \middle| w_{1:N}^{1:P}, b^{1:M}\right) = \mathcal{P}\left(\mathbf{x}_{1:N}^{\mathcal{M}_0} \middle| \text{MSD}\right) \mathcal{P}\left(\text{MSD} \middle| \mathbf{x}_{1:N}^{\mathcal{M}_1}\right) \mathcal{P}\left(\mathbf{x}_{1:N}^{\mathcal{M}_1} \middle| w_{1:N}^{1:P}\right). \tag{S.49}$$

Using the logic of ancestral sampling, we can now sample all trajectories and MSD in three steps:

1. Sample the trajectories of emitters with active loads from $\mathcal{P}\left(\mathbf{x}_{1:N}^{\mathcal{M}_1} \middle| w_{1:N}^{1:P}\right)$;
2. Sampling the MSD from $\mathcal{P}\left(\text{MSD} \middle| \mathbf{x}_{1:N}^{\mathcal{M}_1}\right)$;
3. Sample the trajectories of emitters with inactive loads from $\mathcal{P}\left(\mathbf{x}_{1:N}^{\mathcal{M}_0} \middle| \text{MSD}\right)$.

For mathematical simplicity, we define the diffusion coefficient as $D = \frac{\text{MSD}}{2\tau}$, where $\tau$ is the frame period, in section C.3 only.

##### C.3.1 Active trajectories

The target distribution of active trajectories, denoted by $\mathcal{P}\left(\mathbf{x}_{1:N}^{\mathcal{M}_1} \middle| w_{1:N}^{1:P}\right)$, has not yet been derived. Therefore, we must first derive this distribution. We begin by writing

$$\mathcal{P}\left(\mathbf{x}_{1:N}^{\mathcal{M}_1} \middle| w_{1:N}^{1:P}\right) = \frac{1}{\mathcal{P}\left(w_{1:N}^{1:P}\right)} \mathcal{P}\left(w_{1:N}^{1:P}, \mathbf{x}_{1:N}^{\mathcal{M}_1} \middle| w_{1:N}^{1:P}\right). \tag{S.50}$$

After omitting the evidence term (the normalization), this becomes

$$\mathcal{P}\left(\mathbf{x}_{1:N}^{\mathcal{M}_1} \middle| w_{1:N}^{1:P}\right) \propto \mathcal{P}\left(w_{1:N}^{1:P}, \mathbf{x}_{1:N}^{\mathcal{M}_1} \middle| h, F\right). \tag{S.51}$$

Next, we can write

$$\begin{aligned} \mathcal{P}\left(w_{1:N}^{1:P}, \mathbf{x}_{1:N}^{\mathcal{M}_1} \middle| h, F\right) &\propto \int_0^\infty \mathrm{d}D \int \mathrm{d}\mathbf{x}_{1:N}^{\mathcal{M}_0} \mathcal{P}\left(D, w_{1:N}^{1:P}, \mathbf{x}_{1:N}^{\mathcal{M}_0}, \mathbf{x}_{1:N}^{\mathcal{M}_1} \middle| h, F\right) \\ &\propto \mathcal{P}\left(w_{1:N}^{1:P} \middle| b^{1:M}, \mathbf{x}_{1:N}^{1:M}\right) \int_0^\infty \mathrm{d}D \mathcal{P}(D) \int \mathrm{d}\mathbf{x}_{1:N}^{\mathcal{M}_0} \mathcal{P}\left(\mathbf{x}_{1:N}^{\mathcal{M}_0}, \mathbf{x}_{1:N}^{\mathcal{M}_1} \middle| D\right). \end{aligned} \tag{S.52}$$

To evaluate this double integral, we start by writing

$$\mathcal{P}\left(\mathbf{x}_{1:N}^{\mathcal{M}_0}, \mathbf{x}_{1:N}^{\mathcal{M}_1} \middle| D\right) = \mathcal{P}\left(\mathbf{x}_{1:N}^{\mathcal{M}_0} \middle| D\right) \mathcal{P}\left(\mathbf{x}_{1:N}^{\mathcal{M}_1} \middle| D\right), \tag{S.53}$$

as the trajectories of different emitters are de-correlated. Additionally, $\int \mathrm{d}\mathbf{x}_{1:N}^{\mathcal{M}_0}$ represents the integration of every single element of $\mathbf{x}_{1:N}^{\mathcal{M}_0}$ over the entire real axis. This leads to

$$\int \mathrm{d}\mathbf{x}_{1:N}^{\mathcal{M}_0} \mathcal{P}\left(\mathbf{x}_{1:N}^{\mathcal{M}_0}, \mathbf{x}_{1:N}^{\mathcal{M}_1} \middle| D\right) = \mathcal{P}\left(\mathbf{x}_{1:N}^{\mathcal{M}_1} \middle| D\right) \int \mathrm{d}\mathbf{x}_{1:N}^{\mathcal{M}_0} \mathcal{P}\left(\mathbf{x}_{1:N}^{\mathcal{M}_0} \middle| D\right) = \mathcal{P}\left(\mathbf{x}_{1:N}^{\mathcal{M}_1} \middle| D\right), \tag{S.54}$$

which results in

$$\mathcal{P}\left(\mathbf{x}_{1:N}^{\mathcal{M}_1} \middle| w_{1:N}^{1:P}\right) \propto \mathcal{P}\left(w_{1:N}^{1:P} \middle| b^{1:M}, \mathbf{x}_{1:N}^{1:M}\right) \int_0^\infty \mathrm{d}D \mathcal{P}(D) \mathcal{P}\left(\mathbf{x}_{1:N}^{\mathcal{M}_1} \middle| D\right). \tag{S.55}$$

To continue, we must calculate the probability of active trajectories, $\mathbf{x}_{1:N}^{\mathcal{M}_1}$, given $D$, and specify its prior, $\mathcal{P}(D)$. As previously mentioned in equation (S.19), we consider the Brownian motion model in this study. Therefore, the probability of $\mathbf{x}_{1:N}^{\mathcal{M}_1}$ given $D$ can be expressed as

$$\mathcal{P}\left(\mathbf{x}_{1:N}^{\mathcal{M}_1} \middle| D\right) = \prod_{m \in \mathcal{M}_1} \left[\prod_{n=1}^{N-1} \mathcal{P}\left(\mathbf{x}_{n+1}^m \middle| \mathbf{x}_n^m, D\right)\right] \mathcal{P}\left(\mathbf{x}_1^m\right). \tag{S.56}$$

In this equation, $\mathcal{P}\left(\mathbf{x}_1^m\right)$ is the prior probability distribution for the initial position of the emitter, which is independent of the diffusion coefficient and therefore constant in the integral of equation (S.55). We will address this distribution shortly.

The Brownian motion model also dictates that

$$\mathcal{P}\left(\mathbf{x}_{n+1}^m \middle| \mathbf{x}_n^m, D\right) = \mathbf{Normal}_3\left(\mathbf{x}_{n+1}^m; \mathbf{x}_n^m, 2D\tau\right) \propto D^{-\frac{3}{2}} \exp\left(-\frac{\left|\mathbf{x}_{n+1}^m - \mathbf{x}_n^m\right|^2}{4D\tau}\right).$$

Multiplying the equation above by all emitters with the load being 1 and all time points yields

$$\prod_{m\in\mathcal{M}_1}\prod_{n=1}^{N-1} \mathcal{P}\left(\mathbf{x}_{n+1}^m \middle| \mathbf{x}_n^m, D\right) \propto D^{-\frac{3}{2}B(N-1)} \exp\left(-\frac{1}{4D\tau}\sum_{m\in\mathcal{M}_1}\sum_{n=1}^{N-1}\left|\mathbf{x}_{n+1}^m - \mathbf{x}_n^m\right|^2\right), \tag{S.57}$$

where $B$ is the cardinality (number of elements) of the active load set $\mathcal{M}_1$. This $D$-dependence can actually be represented by an Inverse Gamma distribution of $D$. Put differently, the Inverse Gamma distribution is the conjugate prior to this likelihood. Therefore, we choose an Inverse Gamma distribution, or conjugate prior, for the diffusion coefficient as

$$D \sim \mathbf{InvGamma}\left(\phi_D, \phi_D\chi_D\right). \tag{S.58}$$

By doing so, we obtain the following equation

$$\begin{aligned}&\mathcal{P}\left(D\right)\mathcal{P}\left(\mathbf{x}_{1:N}^{\mathcal{M}_1}\middle| D\right)\\ &\propto \left[\prod_{m\in\mathcal{M}_1}\mathcal{P}\left(\mathbf{x}_1^m\right)\right] D^{-\frac{3}{2}B(N-1)-\phi_D-1} \exp\left(-\frac{\phi_D\chi_D}{D} - \frac{1}{4D\tau}\sum_{m\in\mathcal{M}_1}\sum_{n=1}^{N-1}\left|\mathbf{x}_{n+1}^m - \mathbf{x}_n^m\right|^2\right).\end{aligned} \tag{S.59}$$

By substituting equations (S.58) and (S.59) into the integrand of equation (S.55) and evaluating the integral, we get

$$\begin{aligned}&\int_0^\infty \mathrm{d}D\mathcal{P}\left(\mathbf{x}_{1:N}^{\mathcal{M}_1}\middle| D\right)\mathcal{P}\left(D\right)\\ &\propto \left[\prod_{m\in\mathcal{M}_1}\mathcal{P}\left(\mathbf{x}_1^m\right)\right]\left(\phi_D\chi_D + \frac{1}{4\tau}\sum_{m\in\mathcal{M}_1}\sum_{n=1}^{N-1}\left|\mathbf{x}_{n+1}^m - \mathbf{x}_n^m\right|^2\right)^{-\frac{3}{2}B(N-1)-\phi_D}.\end{aligned} \tag{S.60}$$

Furthermore, we can see from equations (S.16) to (S.18) that the initial positions of all emitters are not correlated. This can be mathematically expressed as

$$\mathcal{P}\left(\mathbf{x}_1^{\mathcal{M}_1}\right) = \prod_{m\in\mathcal{M}_1}\mathbf{Normal}\left(x_{(1)}^m; \mu_{xy}, \sigma_{xy}^2\right)\mathbf{Normal}\left(y_{(1)}^m; \mu_{xy}, \sigma_{xy}^2\right)\mathbf{Normal}\left(z_{(1)}^m; \mu_z, \sigma_z^2\right). \tag{S.61}$$

Using this information, we can derive the target distribution from which we can sample trajectories associated with loads 1.

$$\mathcal{P}\left(\mathbf{x}_{1:N}^{\mathcal{M}_1} \middle| w_{1:N}^{1:P}\right) \\ \propto \mathcal{P}\left(\mathbf{x}_1^{\mathcal{M}_1}\right) \mathcal{P}\left(w_{1:N}^{1:P} \middle| b^{1:M}, \mathbf{x}_{1:N}^{1:M}\right) \left(\phi_D \chi_D + \frac{1}{4\tau} \sum_{m\in\mathcal{M}_1} \sum_{n=1}^{N-1} \left|\mathbf{x}_{n+1}^m - \mathbf{x}_n^m\right|^2\right)^{-\frac{3}{2}B(N-1)-\phi_D}. \tag{S.62}$$

##### C.3.2 MH random walk

Since the fully written-out posterior probability distribution in equation (S.62) is not suitable for direct sampling, we use the MH algorithm to generate new samples. In the proposal distribution, we add a Normal perturbation to the current trajectory to produce proposed trajectories

$$\mathbb{Q}\left(\mathbf{x}_{1:N}^{\mathcal{M}_1,\text{prop}} \middle| \mathbf{x}_{1:N}^{\mathcal{M}_1,\text{old}}\right) = \mathbf{Normal}_3\left(\mathbf{x}_{1:N}^{\mathcal{M}_1,\text{prop}}; \mathbf{x}_{1:N}^{\mathcal{M}_1,\text{old}}, \mathbf{\Sigma_x}\right). \tag{S.63}$$

The perturbation size is controlled by the elements of the diagonal matrix $\mathbf{\Sigma_x}$, which can be manually set or automatically tuned for a target acceptance ratio. This proposal is advantageous because it satisfies the symmetry $\mathbb{Q}\left(\mathbf{x}_{1:N}^{\mathcal{M}_1,\text{prop}} \middle| \mathbf{x}_{1:N}^{\mathcal{M}_1,\text{old}}\right) = \mathbb{Q}\left(\mathbf{x}_{1:N}^{\mathcal{M}_1,\text{old}} \middle| \mathbf{x}_{1:N}^{\mathcal{M}_1,\text{prop}}\right)$, which allows us to simplify the acceptance ratio to

$$r = \min\left\{\frac{\mathcal{P}\left(\mathbf{x}_{1:N}^{\mathcal{M}_1,\text{prop}} \middle| w_{1:N}^{1:P}\right)}{\mathcal{P}\left(\mathbf{x}_{1:N}^{\mathcal{M}_1,\text{old}} \middle| w_{1:N}^{1:P}\right)}, 1\right\}, \tag{S.64}$$

Based on equation (S.62), the ratio between the posterior probabilities can be expressed as

$$\frac{\mathcal{P}\left(\mathbf{x}_{1:N}^{\mathcal{M}_1,\text{prop}} \middle| w_{1:N}^{1:P}\right)}{\mathcal{P}\left(\mathbf{x}_{1:N}^{\mathcal{M}_1,\text{old}} \middle| w_{1:N}^{1:P}\right)} = \frac{\mathcal{L}\left(\mathbf{x}_{1:N}^{\mathcal{M}_1,\text{prop}}\right)}{\mathcal{L}\left(\mathbf{x}_{1:N}^{\mathcal{M}_1,\text{old}}\right)} \frac{\mathcal{P}\left(\mathbf{x}_1^{1:M,\text{prop}}\right)}{\mathcal{P}\left(\mathbf{x}_1^{1:M,\text{old}}\right)} \\ \times \left(\frac{\phi_D \chi_D + \frac{1}{4\tau} \sum_{m\in\mathcal{M}_1} \sum_{n=1}^{N-1} \left|\mathbf{x}_{n+1}^{m,\text{prop}} - \mathbf{x}_n^{m,\text{prop}}\right|^2}{\phi_D \chi_D + \frac{1}{4\tau} \sum_{m\in\mathcal{M}_1} \sum_{n=1}^{N-1} \left|\mathbf{x}_{n+1}^{m,\text{old}} - \mathbf{x}_n^{m,\text{old}}\right|^2}\right)^{-\frac{3}{2}B(N-1)-\phi_D}. \tag{S.65}$$

The acceptance ratio and all probability densities in the implementation of this sampler are calculated in logarithmic space. The terms in equation (S.65) have been defined earlier in equation (S.24), equation (S.64) and equations (S.20) to (S.22), so they will not be further discussed here. Another computational concern when implementing this algorithm is the mixing time, or the time it takes to find the samples with the maximum posterior. In order to achieve optimal performance, it is necessary to find a balance between the acceptance ratio and perturbation size. In particular, this active trajectory sampler allows for partial perturbation, which means that we can perturb segments of one or multiple emitter trajectories in random order while keeping other parts of the trajectories unchanged. We note that this simply means that elements in the matrix $\mathbf{\Sigma}_{\mathbf{x}}$ from equation (S.63) are set to zero, in turn, and this does not affect the mathematical derivations of this section.

##### C.3.3 Sampling diffusion coefficient and inactive trajectories

Similar to the derivations in the previous sections, we begin by defining the target distribution $\mathcal{P}\left(D \middle| \mathbf{x}_{1:N}^{\mathcal{M}_1}\right)$, which can be expressed using Bayes' theorem as

$$\mathcal{P}\left(D \middle| \mathbf{x}_{1:N}^{\mathcal{M}_1}\right) \propto \mathcal{P}\left(\mathbf{x}_{1:N}^{\mathcal{M}_1} \middle| D\right) \mathcal{P}\left(D\right). \tag{S.66}$$

Both terms on the right side of equation (S.66) were previously discussed when calculating the integrand in equation (S.59). Therefore, for the posterior $\mathcal{P}\left(D \middle| \mathbf{x}_{1:N}^{\mathcal{M}_1}\right)$, we write

$$\mathcal{P}\left(D \middle| \mathbf{x}_{1:N}^{\mathcal{M}_1}\right) \propto D^{-\frac{3}{2}B(N-1)-\phi_D-1} \exp\left(-\frac{\phi_D \chi_D}{D} - \frac{1}{4D\tau} \sum_{m\in\mathcal{M}_1} \sum_{n=1}^{N-1} \left|\mathbf{x}_{n+1}^m - \mathbf{x}_n^m\right|^2\right), \tag{S.67}$$

or equivalently

$$D \sim \mathbf{InvGamma}\left(\phi_D + \frac{3}{2}B\left(N-1\right), \phi_D \chi_D + \frac{1}{4\tau} \sum_{m\in\mathcal{M}_1} \sum_{n=1}^{N-1} \left|\mathbf{x}_{n+1}^m - \mathbf{x}_n^m\right|^2\right). \tag{S.68}$$

From this, we can see that the diffusion coefficient can be sampled directly from the resulting posterior probability distribution above.

Finally, we must update the trajectories of emitters with inactive loads. The target distribution in this case is $\mathcal{P}\left(\mathbf{x}_{1:N}^{\mathcal{M}_0} \middle| D\right)$. As mentioned at the beginning of section C.3, these emitters do not contribute to observations and are therefore only sampled based on their prior distribution, defined by equations (S.16) to (S.18) and equation (S.19).

#### C.4 Special trajectory proposals

In section C.3.1, we described how we sample active load trajectories using the MH algorithm with a Normal perturbation proposal distribution. However, because each coordinate of an active load at any time point represents one dimension in the sample space of the target distribution (as shown in equation (S.62)), it can be challenging to achieve rapid mixing using only one type of proposal. Therefore, we have implemented two additional proposal distributions for active load trajectories to improve computational efficiency: the flip proposal and the swap proposal. These "special" proposals take advantage of certain symmetries in our model while the target distribution remains unchanged.

##### C.4.1 Flip proposal

The flip proposal in our model is derived from the PSF form. As shown in equations (S.7) and (S.8), the considered PSF is symmetric about the focal plane ($xy$-plane). This means that flipping an emitter's spatial position about the focal plane, or mathematically,

$$z_n^{m,\text{prop}} = -z_n^m, \; z_n^{m,\text{old}} = z_n^m, \tag{S.69}$$

does not alter the results of equations (S.20) to (S.22), or the observation likelihood in equation (S.24). As a result, the ratio between the target distributions, which is a part of the acceptance ratio, is simplified to

$$\frac{\mathcal{P}\left(\mathbf{x}_{1:N}^{\mathcal{M}_1,\text{prop}} \middle| w_{1:N}^{1:P}, \text{MSD}\right)}{\mathcal{P}\left(\mathbf{x}_{1:N}^{\mathcal{M}_1,\text{old}} \middle| w_{1:N}^{1:P}, \text{MSD}\right)} = \frac{\mathcal{P}\left(\mathbf{x}_{1:N}^{\mathcal{M}_1,\text{prop}} \middle| \text{MSD}\right)}{\mathcal{P}\left(\mathbf{x}_{1:N}^{\mathcal{M}_1,\text{old}} \middle| \text{MSD}\right)}. \tag{S.70}$$

Substituting equation (S.57) into the previous equation yields

$$\frac{\mathcal{P}\left(\mathbf{x}_{1:N}^{\mathcal{M}_1,\text{prop}} \middle| w_{1:N}^{1:P}, \text{MSD}\right)}{\mathcal{P}\left(\mathbf{x}_{1:N}^{\mathcal{M}_1,\text{old}} \middle| w_{1:N}^{1:P}, \text{MSD}\right)} = \exp\left(-\sum_{m\in\mathcal{M}_1}\sum_{n=1}^{N-1}\frac{\left|-\mathbf{x}_{n+1}^{m,\text{prop}} - \mathbf{x}_n^{m,\text{prop}}\right|^2 - \left|\mathbf{x}_{n+1}^{m,\text{old}} - \mathbf{x}_n^{m,\text{old}}\right|^2}{2\text{MSD}}\right). \tag{S.71}$$

In practice, we randomly select one emitter, denoted as $m'$ and flip all of its positions after a randomly chosen time point $n'$. This allows us to further simplify equation (S.71) to

$$\frac{\mathcal{P}\left(\mathbf{x}_{1:N}^{\mathcal{M}_1,\text{prop}} \middle| w_{1:N}^{1:P}, \text{MSD}\right)}{\mathcal{P}\left(\mathbf{x}_{1:N}^{\mathcal{M}_1,\text{old}} \middle| w_{1:N}^{1:P}, \text{MSD}\right)} = \exp\left(-\frac{\left|-z_{n'+1}^{m'} - z_{n'}^{m'}\right|^2 - \left|z_{n'+1}^{m'} - z_{n'}^{m'}\right|^2}{2\text{MSD}}\right). \tag{S.72}$$

As flipping an emitter's positions twice should naturally result in the same position, the proposal should therefore be symmetric

$$\frac{\mathbb{Q}\left(\mathbf{x}_{1:N}^{\mathcal{M}_1,\text{prop}} \middle| \mathbf{x}_{1:N}^{\mathcal{M}_1,\text{old}}\right)}{\mathbb{Q}\left(\mathbf{x}_{1:N}^{\mathcal{M}_1,\text{old}} \middle| \mathbf{x}_{1:N}^{\mathcal{M}_1,\text{prop}}\right)} = 1. \tag{S.73}$$

The logarithmic acceptance ratio is then given by

$$\ln r = \min\left\{\frac{\left|z_{n'+1}^{m'} - z_{n'}^{m'}\right|^2 - \left|-z_{n'+1}^{m'} - z_{n'}^{m'}\right|^2}{2\text{MSD}}, 0\right\}. \tag{S.74}$$

##### C.4.2 Swap proposal

The swapping operation, like the flip proposal, has the advantage of maintaining the invariance of the observation likelihood and being symmetric. To implement this operation, we randomly select $m_1$ and $m_2$, both from the set $\mathcal{M}_1$, and $n = 1, \ldots, N-1$. We then exchange the positions for all subsequent time points according to the following equation

$$\mathbf{x}_{n:N}^{m_1} \to \mathbf{x}_{n:N}^{m_2}, \mathbf{x}_{n:N}^{m_2} \to \mathbf{x}_{n:N}^{m_1}. \tag{S.75}$$

The logarithmic acceptance ratio for this operation is calculated using the following expression

$$\ln r = \min\left\{ \frac{\left|\mathbf{x}_{n+1}^{m_2} - \mathbf{x}_n^{m_2}\right|^2 + \left|\mathbf{x}_{n+1}^{m_1} - \mathbf{x}_n^{m_1}\right|^2 - \left|\mathbf{x}_{n+1}^{m_2} - \mathbf{x}_n^{m_1}\right|^2 - \left|\mathbf{x}_{n+1}^{m_1} - \mathbf{x}_n^{m_2}\right|^2}{2\mathrm{MSD}}, 0 \right\}. \tag{S.76}$$
